# 3D super-resolution microscopy performance and quantitative analysis assessment using DNA-PAINT and DNA origami test samples

**DOI:** 10.1101/626887

**Authors:** Ruisheng Lin, Alexander H. Clowsley, Tobias Lutz, David Baddeley, Christian Soeller

## Abstract

Assessment of the imaging quality in localisation-based super-resolution techniques relies on an accurate characterisation of the imaging setup and analysis procedures. Test samples can provide regular feedback on system performance and facilitate the implementation of new methods. While multiple test samples for regular, 2D imaging are available, they are not common for more specialised imaging modes. Here, we analyse robust test samples for 3D and quantitative super-resolution imaging, which are straightforward to use, are time-and cost-effective and do not require experience beyond basic laboratory and imaging skills. We present two options for assessment of 3D imaging quality, the use of microspheres functionalised for DNA-PAINT and a commercial DNA origami sample. A method to establish and assess a qPAINT workflow for quantitative imaging is demonstrated with a second, commercially available DNA origami sample.

## 1. Introduction

Super-resolution (SR) imaging techniques have been revolutionising optical microscopy and push the achievable resolution to greater than tenfold of that obtainable with diffraction limited conventional microscopy. Single molecule localisation microscopy (SMLM) is one approach to optical SR and encompasses methods such as PALM [1], STORM [2] and DNA-PAINT [3] which rely on the temporal modulation and accumulation of non-overlapping single-molecule events to gradually construct an image. Applications of SMLM in visualising complex cellular structures at the nanoscale span across various areas in biological and medical research, such as studies of microtubules [4], eukaryotes [5], nuclear biology [6] and cardiac myocytes [7, 8], for a review see [9].

In general, in microscopy it is increasingly appreciated that robust test procedures to evaluate system performance are essential since both alignment issues and component performance can degrade instrument performance below design specification [10–13]. At least of equal importance, such performance checks enable the comparison of data obtained in different laboratories with greater confidence. The procedures generally rely on the availability of suitable test samples that are often specifically constructed to enable reproducible implementation and interpretation of test procedures [14, 15].

In SMLM systems performance is typically limited by at least three factors: (1) the labelling density and overall quality of the sample [16–19]; (2) the photon yield associated with each single-molecule event for localisation precision [20–24]; (3) the stabilisation of the system against mechanical and thermal drift over long acquisition times. In practice, it is therefore essential to characterise an SMLM system and optimise its performance prior to embarking on a new project or one involving complex biological samples.

The testing of super-resolution imaging systems has been discussed previously, e.g. [25–27], and a number of strategies have been developed that rely, for example, on nano-structured test samples [25–32] and software packages that include test and calibration procedures [33]. In addition, due to the importance of the data analysis pipeline in SMLM, software packages for super-resolution have been tested and compared in benchmarking exercises [34, 35].

Here we present a number of robust test and validation procedures for a localisation based super-resolution microscopy system. The samples are based on the DNA-PAINT technique and can be used on various SMLM platforms. First, we perform high-resolution 3D imaging using DNA-PAINT microspheres and commercially available origami test samples. We also demonstrate procedures to test and validate quantitative analysis of DNA-PAINT using information extracted from the localisation data implementing the approach termed qPAINT [8, 36]. We conclude that the advantages of providing clear structures, “best-case-scenario” localisation data with high signal-to-noise ratio, and the comparatively long durability of suitable samples make these procedures and the associated DNA-PAINT based samples convenient off-the-shelf solutions for evaluating and validating the performance of SMLM systems.

## 2. Basic Layout of the SMLM setup

All test and validation procedures were carried out with a custom built microscope setup that was constructed around a conventional widefield fluorescence microscope (Nikon Eclipse Ti-E). A 647 nm diode laser was used to excite ATTO 655 dye molecules in the sample through a high NA TIRF objective. Fluorescence was collected by the same high NA objective and propagated through a Cy5 filter set which matched the emission spectrum of the dye. For 2D super-resolution imaging, an sCMOS camera was installed directly at the microscope side port and aligned with a conjugate sample plane, marked as ‘intermediate image plane’ in Fig. 1. When performing 3D localisation imaging using the biplane approach [37, 38] (section 4), an optical splitter was inserted to equally split the signal between two channels, one of which was re-focused by a very weak (f = 4 m) lens to introduce an axial focus shift (Fig. 1). Alternatively, a phase ramp slider was inserted in the infinity space below the objective lens to implement phase-ramp imaging localisation microscopy (PRILM) for 3D localisation [39]. Hardware control, single molecule localisation and post-processing was performed with a custom, open-source software package called PYME (Python Microscopy Environment). Full details of the setup, software and procedures are provided in section 7.

**Figure 1.**
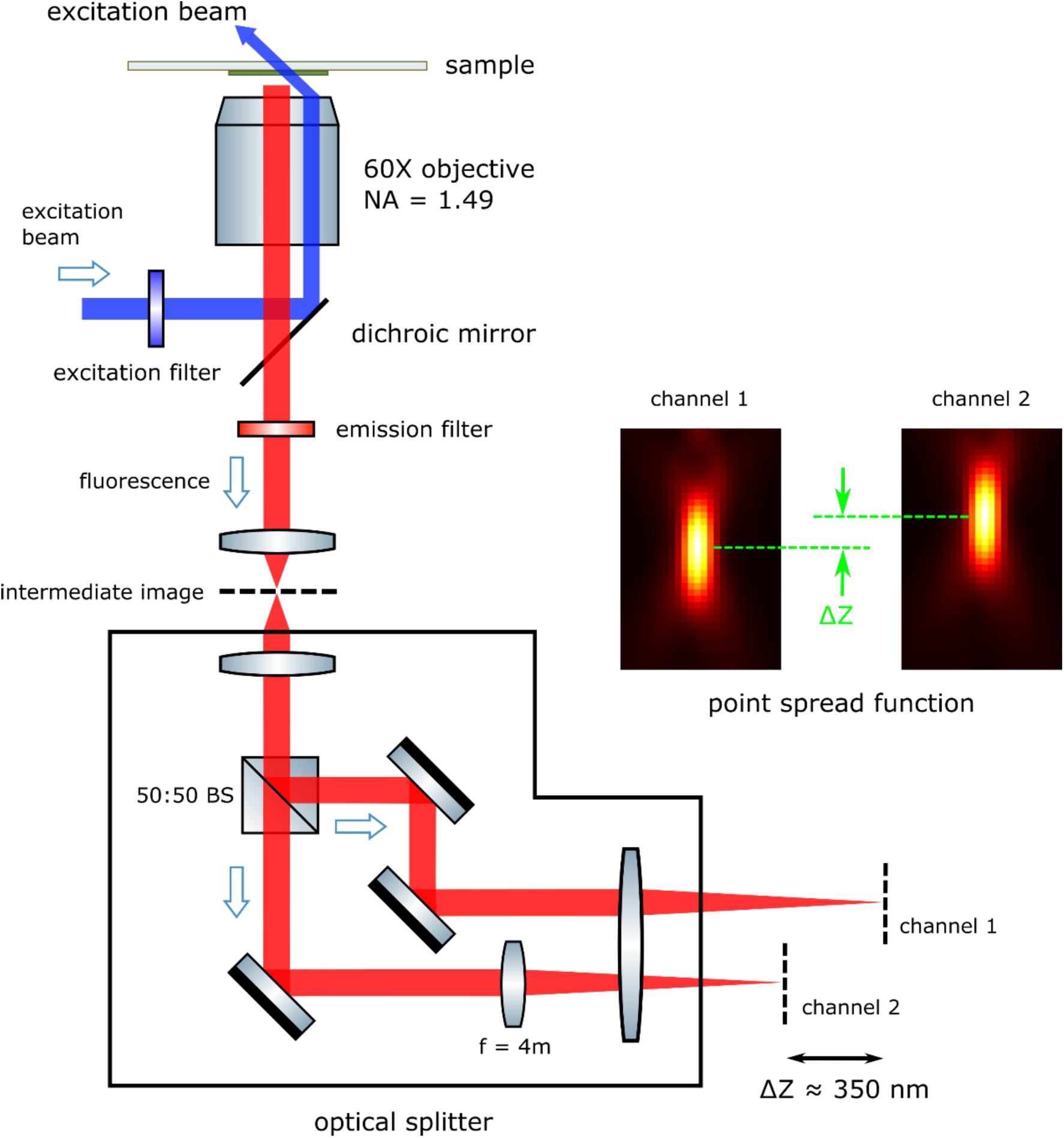
Schematic of a 3D single molecule localisation microscope with biplane 3D localisation. The laser beam is coupled into a conventional fluorescence microscope in TIRF or HILO mode to provide widefield excitation. For biplane 3D localisation an optical splitter is installed between the microscope and the camera. The fluorescence is divided by a 50:50 beam splitter which produces two separate images on both halves of the camera sensor. An f = 4 m lens is inserted into one channel of the splitter device to introduce ∼350 nm shift (approximately half of the axial full-width-half-maximum of the point spread function) between the focus planes.

## 3. Principles of DNA-PAINT and DNA-PAINT based test samples

DNA-PAINT (Point Accumulation Imaging in Nanoscale Topography) [40] is based on the transient binding of fluorescently labelled oligonucleotides [3, 41]. Fluorescently modified strands (“imagers”) in solution are detected only as a diffuse fluorescence background signal, they appear as diffraction-limited spots in camera-frames once they are immobilised upon binding to stationary complementary target strands (“docking strands”) within the sample. For effective localisation based imaging, imager and docking strands are designed so that their average binding time is comparable to the camera frame integration time (typically using frame integration times on the order of 0.1 s). The bright apparent “blinks” that result from the transient immobilisation can be localised with up-to single-nanometre precision, depending on photon yields and background intensities. The diffuse background arising from rapidly diffusing imagers in solution can be minimised by imaging in total internal reflection fluorescence (TIRF) [42] or highly inclined and laminated optical sheet (HILO) [43] imaging modes. In addition to the DNA strand designs, binding times can be further adjusted by modifying buffer conditions. The DNA-PAINT approach enables imaging with high specificity and contrast and, unlike several other localisation based super-resolution techniques, dye photobleaching is effectively negligible because fresh imagers diffuse in from the bulk medium to replace imagers that may have been bleached. This latter point in particular makes DNA-PAINT based samples a very convenient choice, since test samples can be maintained in good condition for weeks once made and, as we show below, in some cases well-sealed samples can remain stable for many months.

The use of samples of known structure is often a pre-requisite for test procedures of super-resolution imaging performance. For this purpose the use of DNA origami [44], i.e. synthetic 2D and 3D DNA structures with feature sizes in the nanometre range, has been proposed and demonstrated by a number of groups [25, 26, 28–31]. For use in DNA-PAINT, DNA origami is especially convenient since DNA strands acting as docking sites can be placed with high precision into DNA origami structures, so that imagers bind to sites that have well defined spacing in the range of <10 nm to >100 nm, depending on origami design and application purpose. Accordingly, the term DNA “nanorulers” has been coined for such preparations [25, 30].

## 4. Assessment of 3D imaging with a DNA-PAINT microsphere sample and a commercial DNA origami sample

Until recently, 3D test samples with well-known 3D structure for super-resolution performance evaluation were not straightforward to obtain. Now, commercial slides with 3D DNA-origami structures are available which provide a stringent test of 3D imaging performance. In addition, we [45] and others [46, 47] have established DNA-PAINT samples based on surface coated microspheres which can be used for a variety of testing purposes associated with DNA-PAINT, including their use as convenient 3D structures of known shape.

### 4.1. 3D Biplane imaging with polystyrene microspheres

Assembly of microsphere-based DNA-PAINT test slides is straightforward, time-and cost-effective (section 7, “Microsphere Sample Preparation for DNA-PAINT”). To validate 3D SMLM imaging, non-fluorescent, 500 nm diameter streptavidin-coated polystyrene microspheres were used as 3D targets with an expected geometry. Using the biplane configuration, images were recorded with the system shown in Fig. 1 with the optical splitting device installed. A 50:50 non-polarising beam splitter cube was inserted to equally divide the image into two channels with one refocused by a weak lens to produce a focal shift of approximately half of the axial full-width-half-maximum of the point spread function.

Prior to image acquisition, remaining sub-diffraction distortions between the two arms of the optical splitter were corrected to a tolerance of ≤10 nm (see section 7, “Calibration of Optical Splitter”) [48]. The biplane configuration point spread function was recorded (see section 7, “Image Acquisition”) by using sparsely distributed fluorescent beads in an aqueous medium, i.e. using the same refractive index as the DNA-PAINT samples. A 647 nm laser was used to excite both the ATTO 655 fluorophore modified imager strands, and the 100 nm red beadsimmobilised on the sample surface serving as fiducial markers (section 7, “Microsphere Sample Preparation for DNA-PAINT”). Illumination was configured in TIRF mode with the evanescent field covering approximately the bottom half of the 500 nm microspheres (Fig. 2A). Sample drift was monitored by recording brightfield images of the microspheres themselves (see section 7, “Experimental Setup”). In addition, images of the fiducial beads were also used for drift correction. We observed that both schemes achieved similar results, however the fiducial correction was slightly more robust and made it the preferred approach when appropriate fiducial markers were present within the sample.

**Figure 2.**
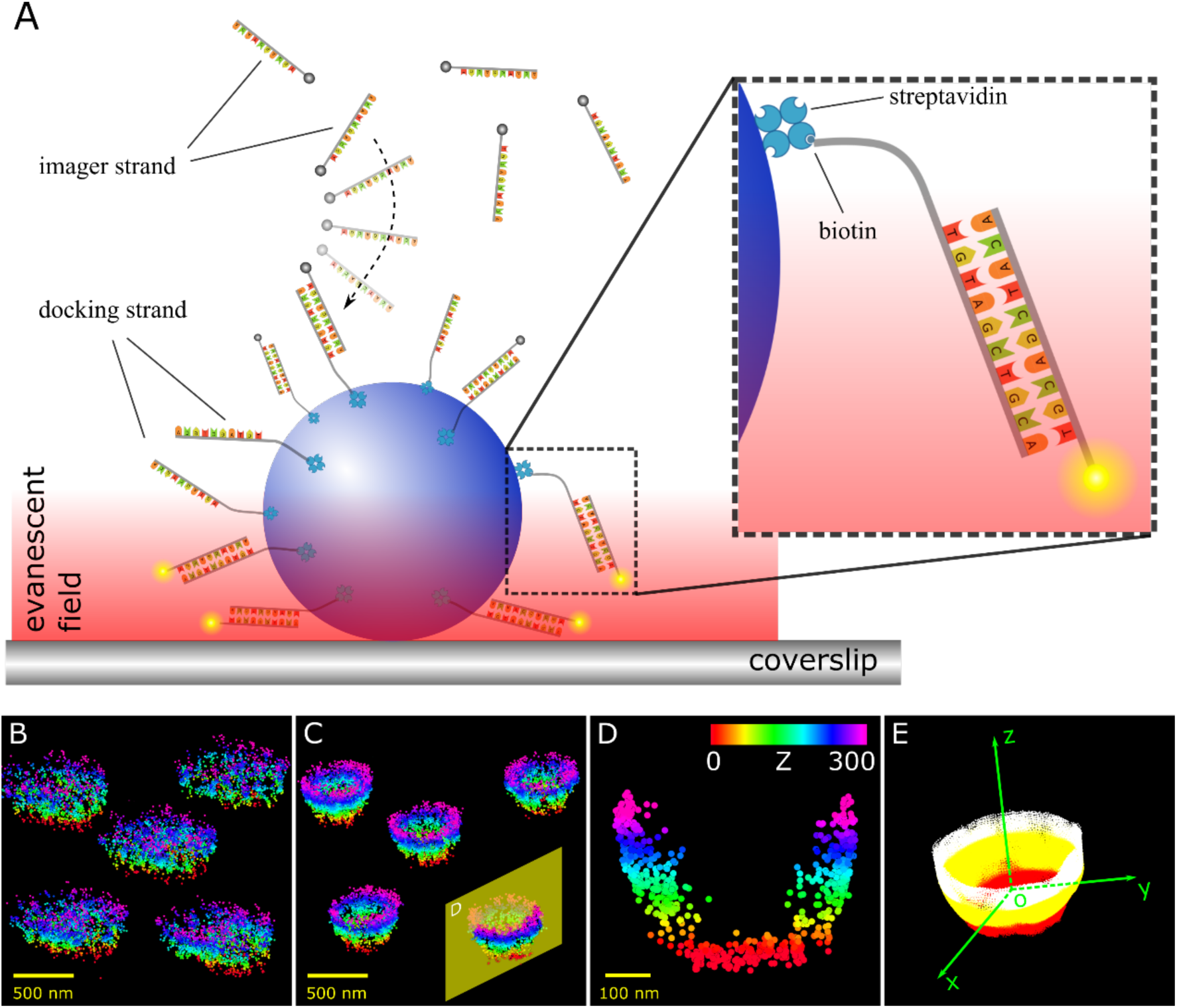
3D DNA-PAINT image of 500 nm polystyrene microspheres using the biplane method. (A) Biotinylated DNA docking strands were attached to the microspheres via streptavidin. Complementary fluorescently modified DNA imager strands were added and left to diffuse freely in the solution. Under laser excitation (TIRF mode), stochastic and transient binding between the docking and imager strands appeared as fluorescent blinks (see inset). (B) Localisation results before 3D drift correction. (C) After 3D drift correction, a hollow spherical structure was observed for each microsphere due to limited thickness of illumination (∼300 nm). (D) The cross-section of a single microsphere in (C) with axial position represented by different colours. (E) An enlarged view of the reconstructed image of a single microsphere in surface plot.

Detected fluorescent blinks were localised using the biplane type 3D localisation algorithm (see section 7, “3D localisation”) which is included in PYME [49]. A set of filtering criteria, such as brightness, lateral and axial localisation precision, were applied to refine the set of localisations for further analysis. Applying fiducial drift correction on the raw localisation clouds (Fig. 2B) yielded a hemispheric structure on each microsphere (Fig. 2C), which was expected due to limited excitation depth. A plot of the cross-section of one of the microspheres exhibited an obvious “U” profile with an axial range of ∼300 nm (Fig. 2D). Reconstructed images of the microsphere using surface plot revealed a clear seamless “cap” structure with a hollow interior (Fig. 2E). Estimation of the localisation precisions, which were ∼10 nm laterally and ∼20 nm axially, suggested that resolutions (full-width-half-maximum) of up to ∼24 nm in the lateral direction and ∼47 nm axially are achievable.

Fig. 3 shows results obtained using the same microsphere sample but using phase ramp 3D imaging (PRILM) [39]. In this method, the PSF was engineered by a phase mask placed in the infinity space behind the objective, producing two lobes whose relative linear orientation is dependent on the axial position. In response to defocus, the two lobes move in opposite directions (Fig. 3A). The use of the phase mask obviates the need of the optical splitter and allows signal to be recorded in one channel. The results (Fig. 3B and Fig. 3C) exhibited good agreement with the biplane approach (Fig. 2).

**Figure 3.**
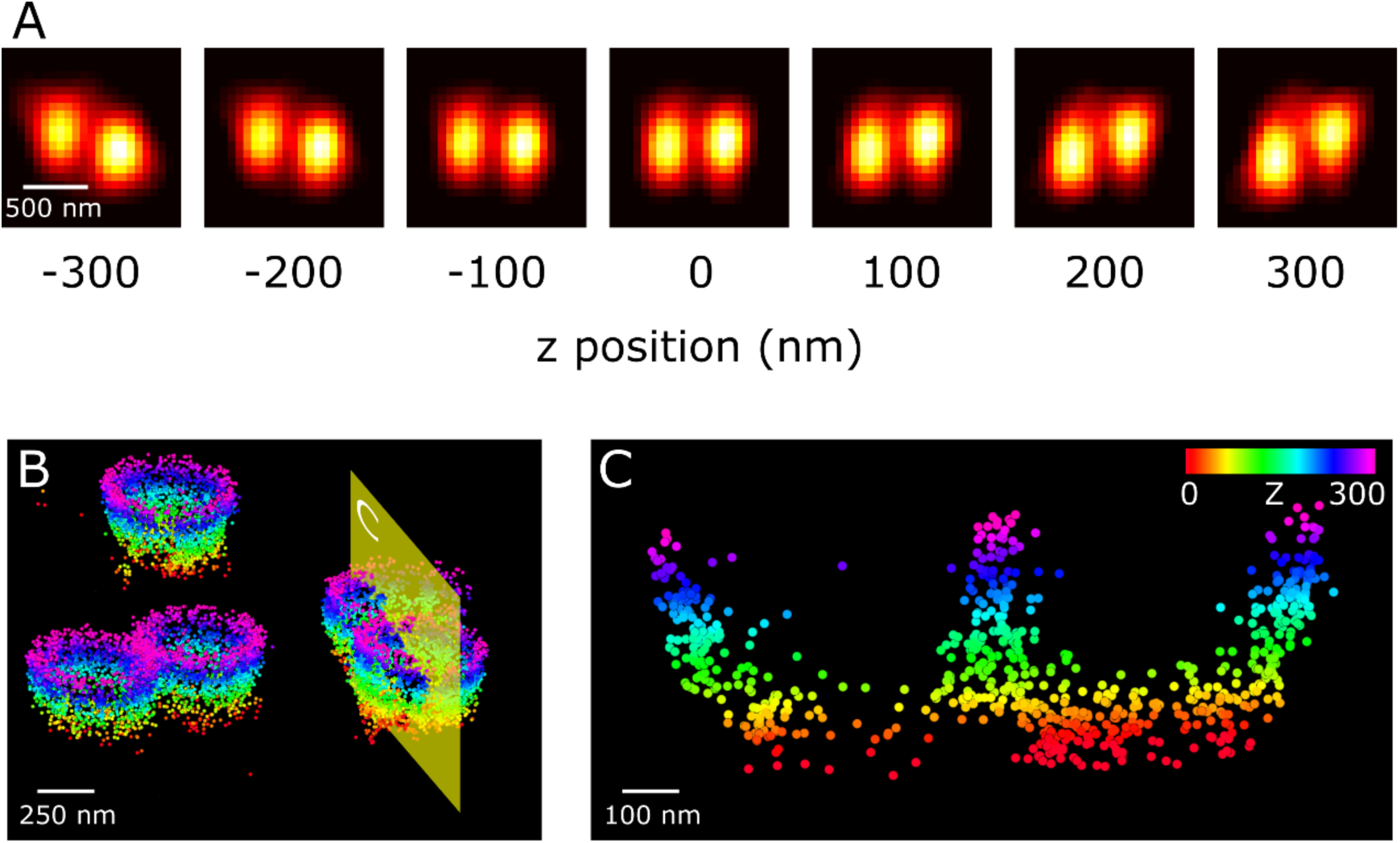
3D Localisation of DNA-PAINT docking strands on the surface of microspheres using the phase ramp method. (A) The measured PSF which is split into two lobes and each lobe moves in opposite directions when changing the axial position. (B) Similar to the biplane mode, after 3D drift correction the localisation cloud of each microsphere exhibits a spherical structure. (C) The cross-section of two adjacent microspheres in (B) shows a clear “UU” shape profile, with z-position ranging from 0 to ∼300 nm.

Error estimation was implemented by utilising a shape accuracy metric. An ellipsoid model was used to fit a localisation data set of a single microsphere acquired using biplane axial localisation, containing ∼3300 data points (Fig. 4A). The fit to the surface of a prolate spheroid (red wireframe) generated best fit estimates of the radius in the equatorial plane as ∼248 nm, which is consistent with the nominal radius of the microspheres (250 nm). In addition, the model also estimates an axial foreshortening factor of 0.65, suggesting that measured z-positions are 1/0.65≈1.5 times larger than the actual z-positions. This axial foreshortening effect, caused by refractive index mismatch between the aqueous solution and oil immersion, should be corrected to obtain the true 3D structure of the specimen (see section 7, “Axial correction of refractive index mismatch”).

**Figure 4.**
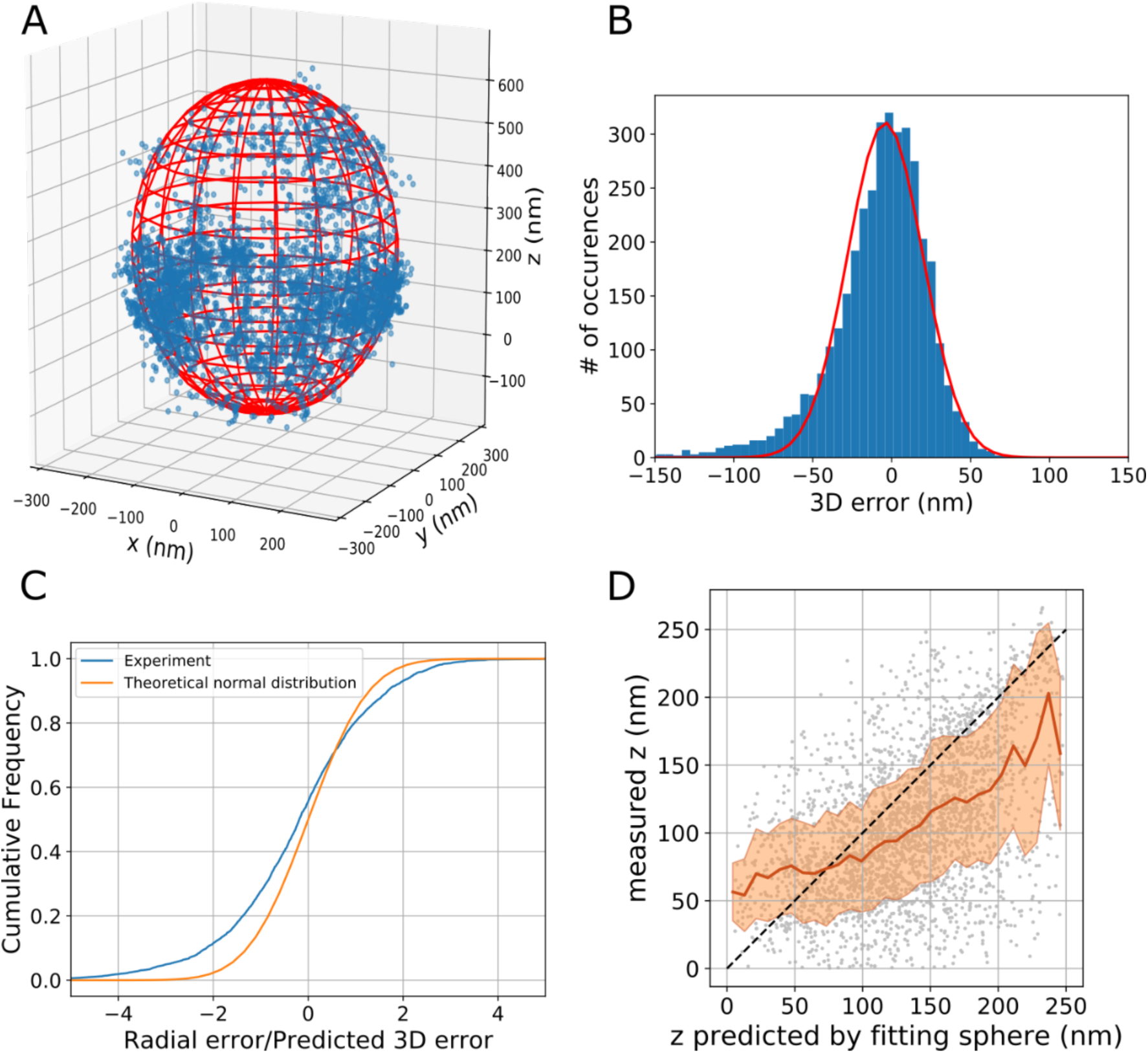
Assessment of 3D localisation using DNA-PAINT data of a 500 nm microsphere. (A) Estimation of the localisation precision using a shape accuracy metric. An ellipsoid model was used to fit the localisation cloud of the microsphere which contains ∼3300 data points. The fit to the surface of prolate spheroid (red wireframe) generated best fit estimates of a radius of the equatorial plane as ∼248 nm and an axial foreshortening factor of 0.65. (B) A histogram plot of the measured radial error, fitted by a Gaussian model (red) with a mean of −4.1 nm and a standard deviation of 23.8 nm. (C) Comparison of the cumulative frequency of the experimental error ratio versus the theoretically expected normal distribution. The difference of the two curves indicates that there is some residual systematic error, such as residual sample drift and non-uniform foreshortening, in the localisation data. (D) Scatter plot of the measured, foreshortening corrected z-positions versus the z-positions predicted by the best-fit model (grey dots). The solid brown curve and shaded area represent z-dependent mean values and standard deviations, respectively.

The overall 3D error reflects contributions both from localisation errors and any additional systematic errors. To estimate the 3D error we calculated the radial deviation of the localisation data points from the best fitting model. A histogram plot of the 3D error is shown in Fig. 4B. A Gaussian curve (red) fitted to the histogram profile indicates a mean value of −4.1 nm with a standard deviation of 23.8 nm.

Systematic errors, such as residual sample drift or distortion of the PSF, were assessed by comparing the radial error with the 3D localisation error predicted by the weighted least squares localisation algorithm [31] at convergence. In the absence of systematic errors, the radial error should consist of only localisation error and their ratio should follow a normal distribution with unit standard deviation. The contribution of systematic errors can be visualised by plotting the cumulative frequency (Fig. 4C) of the experimentally obtained ratio (blue) and a theoretically expected normal distribution (orange). The difference of the two curves indicates a contribution of additional systematic error. In addition, a horizontal shift of the experimental error ratio was observed, reflecting the asymmetric distribution of the overall 3D error.

Fig. 4D shows a scatter plot of the measured, foreshortening corrected z-positions versus the z-positions predicted by the fitting model (grey dots). The z-dependent mean values and their standard deviations are plotted as the solid brown curve and shaded area, respectively. The general trend indicates that the measured z follows the predicted shape albeit with some systematic deviation that may indicate several effects. It could indicate uncorrected drift which could be tested if fiducial markers were present in the sample. In our experiments, fiducial inspection suggested that drift was well corrected. The residual errors indicate that complex shapes in real samples may incur some 3D distortion that is depth dependent. Overall, however, the general 3D topology and local distances were well reconstructed and validated the ability to perform high-resolution 3D imaging.

### 4.2. 3D Biplane imaging with a commercial 3D DNA origami sample

The commercial DNA-origami slide, GATTA-PAINT 3D HiRes 80R (GATTAquant), contains rod-like structures having two sets of docking locations spaced 80.6±20.5 nm apart [26, 50]. These ‘nanopillars’ are anchored in a fashion so that across the sample a proportion are axially positioned approximately vertically with respect to the sample surface, Fig. 5A.

**Figure 5.**
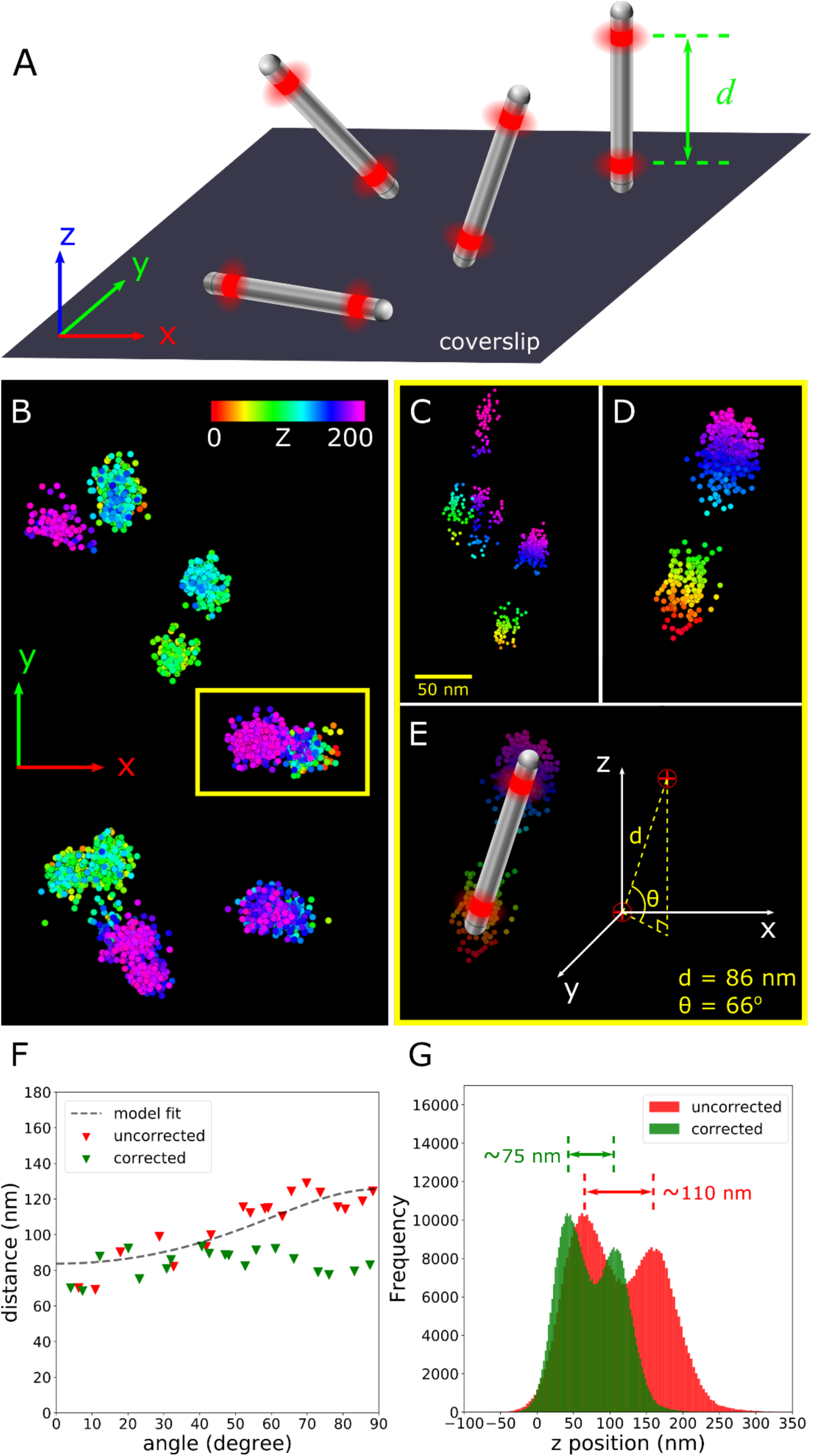
3D DNA-PAINT image of a GATTA-PAINT 3D HiRes 80R test sample using the biplane method. (A) Schematic diagram of the test sample. Nanopillars with two spots separated by a distance *d* show a broad angular distribution in standing, tilted and lying orientations. (B) 2D view of event localisation clouds in the x-y plane obtained by imaging a small section of the 3D test slide. The axial position is coded with a colour map. (C) 3D view of the localisation clouds in the yellow box in (B) without any drift correction. (D) The same localisation clouds in (C) with 3D drift correction. The two spots are well separated. (E) Measurement of the centres of the two clouds with axial foreshortening due to refractive index mismatch corrected shows the distance of the spots and the angle of the nanopillar as 86 nm and 66°. (F) Applying the axial foreshortening correction on 20 randomly selected nanorulers produces an angle-independent length of ∼83.8 nm. (G) Histogram plot of the axial positions shows two clear peaks separated by ∼110 nm (uncorrected, red), or ∼75 nm after correcting foreshortening (green).

The same fluorophore as used in DNA-PAINT microsphere imaging, ATTO 655, was attached to the imager strands. Imaging was performed using the same system shown in Fig. 1 and using the same imaging procedure, except that brighfield tracking was used for drift correction as the sample was manufactured in a sealed slider with no fiducial markers added.

Fig. 5B shows a lateral view of the event localisation data with drift correction applied using the 3D origami sample. Events are coded with colours representing different axial positions. Distance and angular analysis of the nanopillars was performed by fitting a Gaussian mixture model (GMM) [51] to the two drift-corrected data clouds of a single nanoruler. Similar to the analysis for the microsphere, an axial correction was applied to eliminate the foreshortening effect induced by refractive index mismatch [52] (Fig. 5C-E). The correction factor was obtained by fitting 20 randomly selected nanorulers to the model described in [26] (see section 7, “Axial correction of refractive index mismatch”). The fit yielded a nanoruler length of 83.8 nm and an axial foreshortening factor of 0.67, in good agreement with the factor obtained from the microsphere sample. Applying a model that takes axial foreshortening into account produced distance estimates for all the nanorulers in a narrow range around ∼84 nm, regardless of angle (Fig. 5F). A statistical analysis of the distance of all nanopillar sample events within the field of view projected along the z-axis produced a histogram with an apparent separation of the two peaks at ∼110 nm (red). This was reduced to ∼75 nm (green) once the foreshortening factor of 0.67 was taken into account, compatible with the sample specifications [50] provided by the manufacturer (Fig. 5G).

## 5 Establishing qPAINT analysis using a commercial DNA origami sample

Whilst for some applications a significantly enhanced resolution may be sufficient to obtain the structural details of the specimen for qualitative research, the quantitative analysis process qPAINT extracts useful information from localisation data. Procedures to implement qPAINT on highly-resolved super-resolution data has been less widely reported. Here, after establishing the performance of our optical system we repurpose a commercial test slide (GATTA-PAINT HiRes 40R nanorulers, GATTAquant) and show how it can be used to establish and validate the workflow for qPAINT analysis that applies to the DNA-PAINT modality [36]. The slide contains two types of structures [53], which can be compared by qPAINT: (1) a DNA origami structure with docking strands forming trimer structures with a linear spacing of 40 nm, ideal for testing 2D localisation precision and drift correction; (2) a more sparsely distributed larger DNA origami structure used as fiducial markers provides signal for sample tracking and drift correction. Both DNA origami structures use the same type of docking strands, albeit at different quantities. The provided sample is fully sealed and in our hands has given reproducible results for >18 months. Here we use a sample with imagers containing the ATTO 655 fluorochrome.

Fig. 6 shows a super-resolution image that was obtained using the GATTA-PAINT HiRes 40R sample, using a sequence of 70k camera frames at a frame rate of 100 ms/frame. Sample drift was corrected by transmitted light imaging correlation based sample tracking (see section 7, “Experimental Setup”). The rendered super-resolution image is shown in Fig. 6. Both the dense trimer nanorulers (showing ∼40 nm spacing) and the sparser fiducial markers of ∼50 nm diameter were well resolved (Fig. 6A) and demonstrate the use of the sample for lateral resolution checking. For comparison, a diffraction-limited image of the same region (Fig. 6B) was generated by simulation using a Gaussian PSF model, as recording a widefield image is impractical due to the fact that fluorochromes were only attached to the imagers which diffuse freely in the solution. In practice, an additional fluorochrome with different emission spectrum from the imager, for example, a Cy3 dye, can be appended to each docking site to produce continuous fluorescence for labelling checks under widefield illumination [45]. The drift-corrected cloud of localisation (Fig. 6C) shows considerably higher number of detected events on the fiducial marker, an indication that more docking strands are available in these structures.

**Figure 6.**
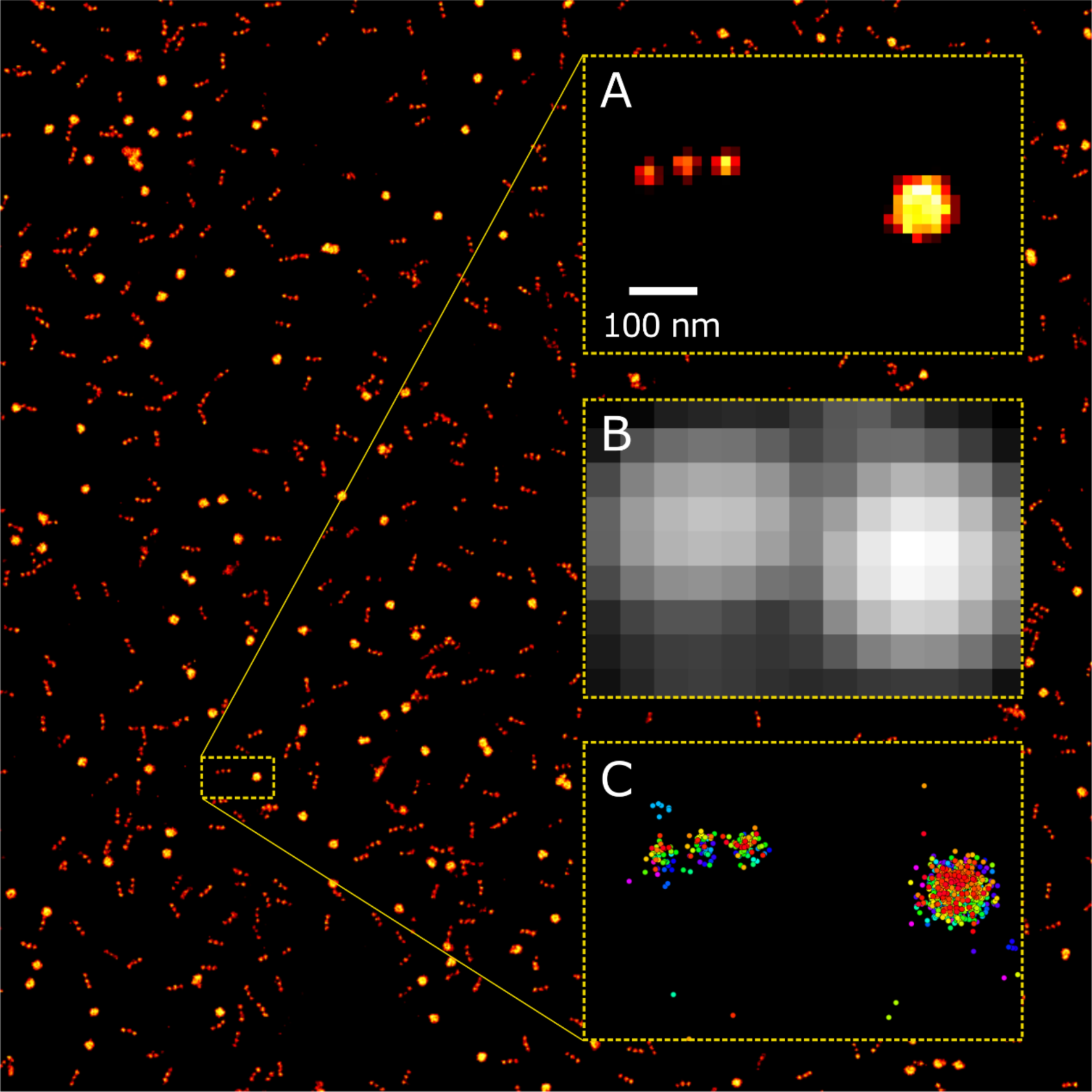
A DNA-PAINT super-resolution image of a GATTA-PAINT HiRes 40R test sample. The three core units, that make up the ∼120 nm nanorulers are well resolved and were widely distributed throughout the sample. Large fiducial markers exist at a lower density amongst the sample. (A) One of the trimer nanorulers with peaks spaced at intervals of 40 nm (centre to centre) and a large fiducial marker. (B) A calculated widefield image of the nanorulers clearly emphasises the diffraction limited nature of the image and demonstrates the inability to resolve it with conventional fluorescence microscopy. (C) Cloud of localisation events of the structures rendered in (A).

The recorded DNA-PAINT data was further processed for quantitative analysis by the qPAINT method. qPAINT utilises an algorithm to quantify the number of binding targets by analysing the temporal properties of DNA-PAINT data, the concept is illustrated in Fig. 7. qPAINT exploits the fact that from a statistical point of view, the number of docking sites, at a fixed imager concentration, is inversely proportional to the mean time between binding events given that the single-molecule DNA hybridisation follows a second order binding kinetics model as described in [36]. The mean time between binding events can be determined by analysing the distribution of dark times between imager-docking hybridisation events appearing as bright blinks in a given region.

**Figure 7.**
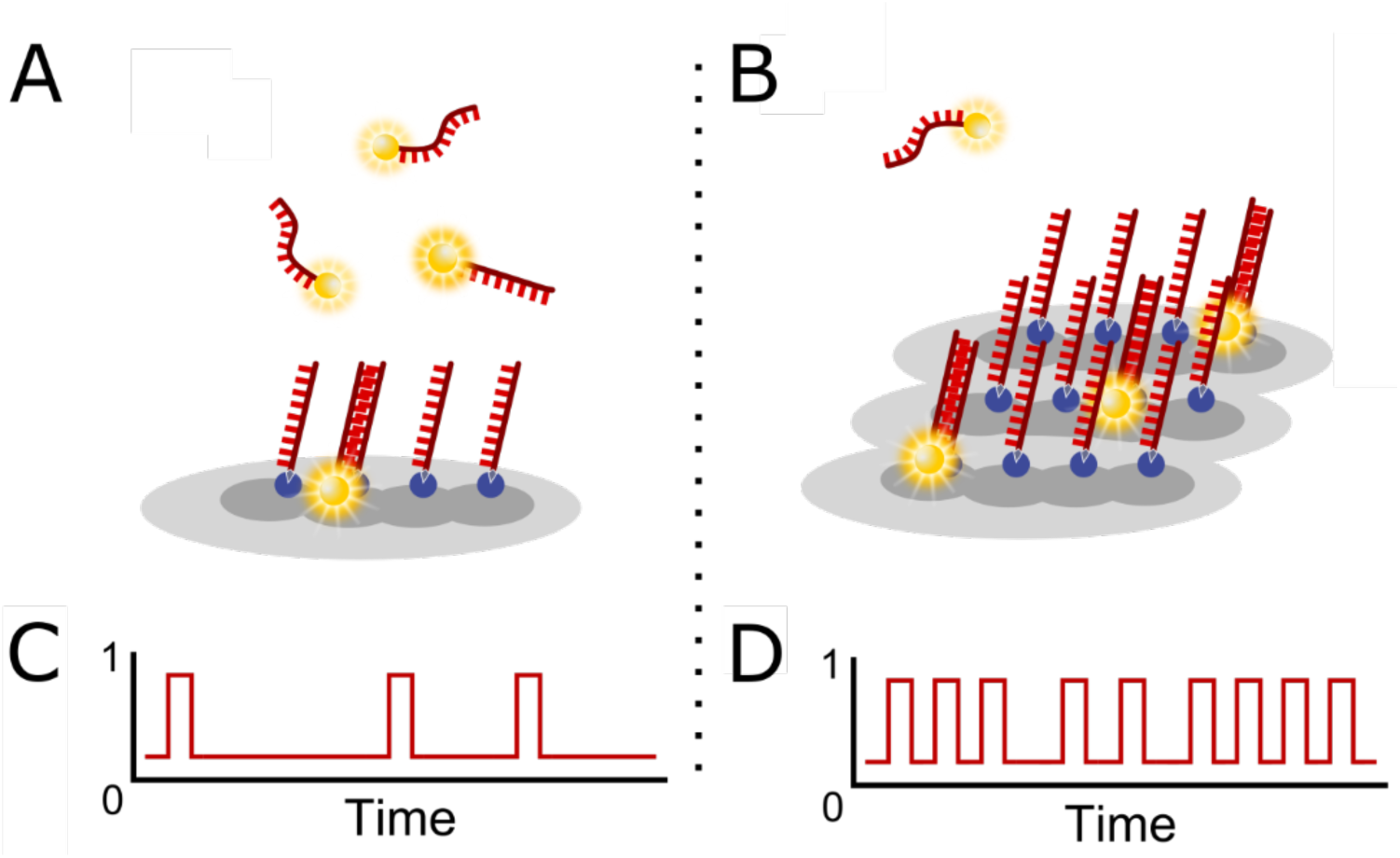
The concept of qPAINT. (A) Imager strands freely diffuse around the sample and transiently bind to complementary fixed docking stations. (B) A target site with three times the amount of docking strands and containing the same imager concentration. (C) & (D) The time traces for detected events in a set time for the scenarios in (A) & (B), respectively.

There are two approaches that can be used to determine the mean dark time in DNA-PAINT experiments, which give essentially equivalent results (Fig. 8A-C). The first one uses fluorescence thresholding by investigating the raw intensity traces versus acquisition time (i.e. frame number) in a selected region of interest. Care needs to be taken to account for drift as structures may move with respect to a fixed region of interest. The second and more convenient method uses detected imager binding events used for generating the super-resolution image. In addition, drift correction, which typically is part of the super-resolution image analysis workflow, ensures that data can be uniquely associated for qPAINT analysis with a single structure. The mean dark time can be measured by fitting a curve to the histogram of dark times determined from the data. If normalised cumulative histograms are used, the expected distribution is an exponential 1-exp (-t/τ_D_) where the mean dark time *τ*_*D*_ is the only free parameter. The relationship between inverse dark times and actual docking site numbers generally requires calibration [8, 36], see also “Quantitative Analysis with qPAINT” in section 7. Here, we determine the relative ratio of the number of accessible docking strands in the nanoruler structure versus the round fiducial structure. We work directly with the inverse of the dark time 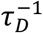 which we call the qPAINT index q_i_ (or qIndex for short) of the region, we typically scaled it by a factor of 100 to obtain numbers of order unity [8].

**Figure 8.**
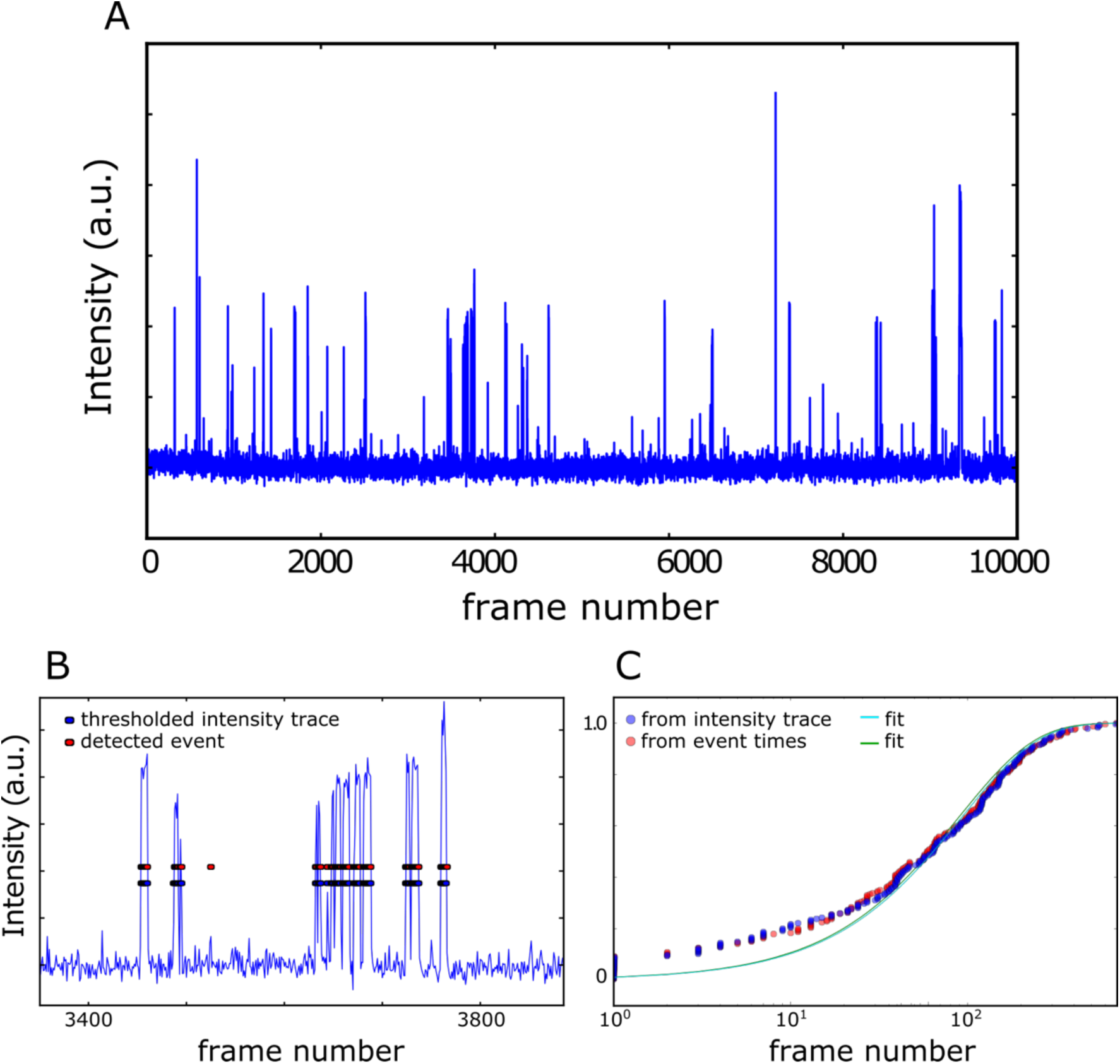
(A) A fluorescent intensity time trace that examines a region of interest containing one of the trimer nanorulers, obtained over the acquisition period of a typical DNA-PAINT experiment. (B) Inspecting an area of ∼400 frames taken from the intensity time plot in (A) exhibits the two approaches to event detection; intensity peaks that surpass a certain threshold or localisation based event detections. (C) Cumulative frequency distributions constructed from dark times obtained by the two ways of analysis show strong agreement. Thresholding (blue circles) and localised event detection (red circles).

Using the qPAINT index terminology, the ratio of docking strands in the nanoruler vs the ratio of docking strands in the fiducial structure can therefore be obtained by the ratio *q*_*i,n*_/*q*_*i,f*_ with those being the estimates of their respective qPAINT indices.

To effectively determine estimates for *q*_*i,n*_ and *q*_*i,f*_ it is necessary to obtain event groups associated with these two types of structures, obtain their mean dark times from the event groups and in this way obtain best estimates of the respective qPAINT indices.

There are a number of ways in which events can be associated with a structure. For example, by directly working with event coordinates, groups of events in ‘clusters’ could be identified [54]. Here we demonstrate an approach that first finds objects by intensity thresholding of the rendered super-resolution image, then we associate groups of events with these image objects and finally select those image objects that correspond to a single nanoruler or the fiducial structures, based on the geometrical properties of these image objects. The procedure is illustrated in Figs. 9-11.

**Figure 9.**
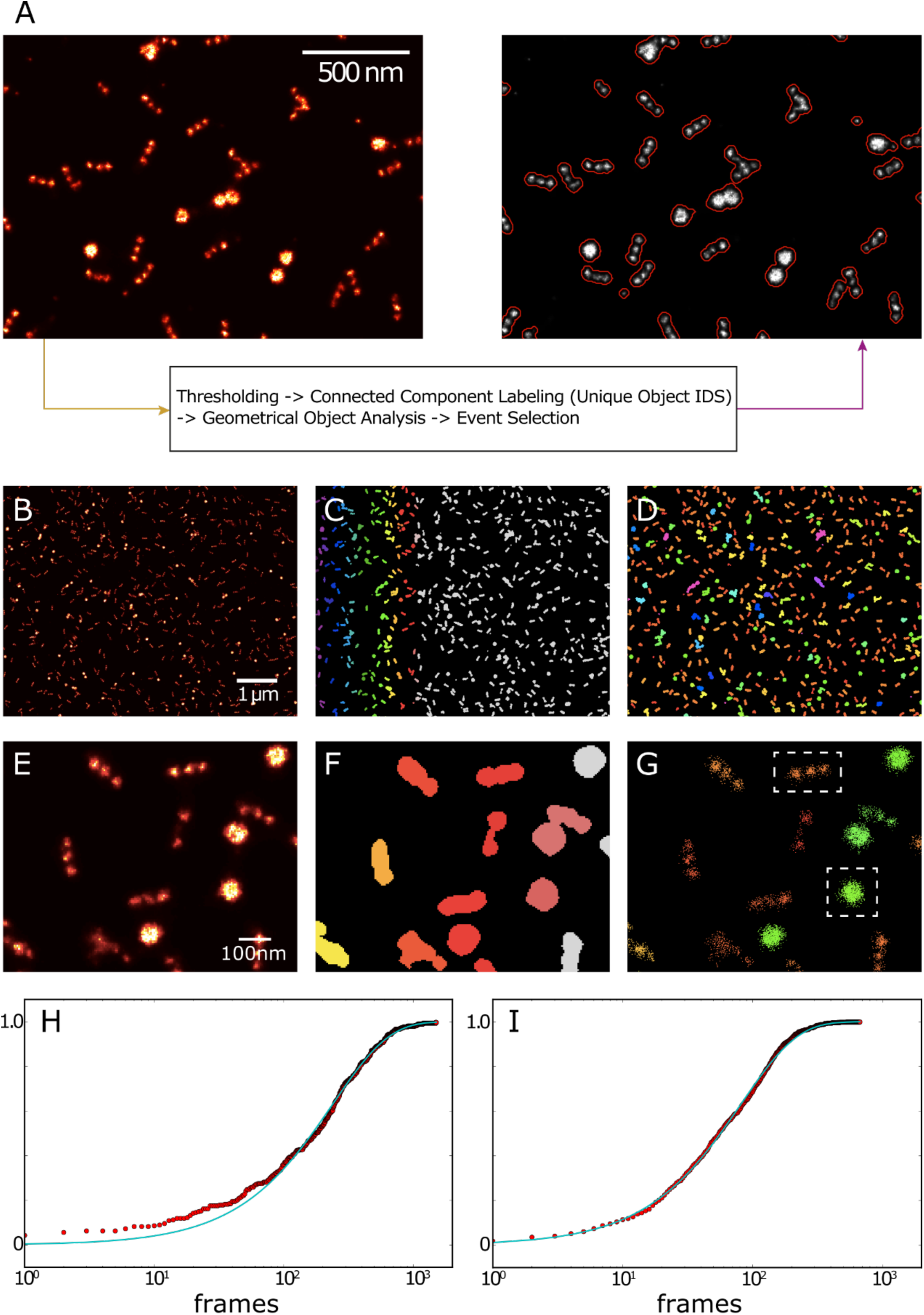
Analysis workflow applied to super-resolution data from a DNA-origami nanoruler. (A) A fractional threshold was applied that kept 80% of the total labelling fraction, above background levels, and a 2D mask of the remaining objects created. Connected components were then assigned individual object identities and their geometrical measurements extracted. Events could then be selected based on a series of filters. An overview of an entire DNA-PAINT rendered image (B), displaying both the nanorulers and fiducial markers. Using the unique object identities, post processing enables the labels to be displayed via a colourmap (C). These labels maintain the object’s geometrical measurements and therefore allow them to be colour-coded based on size (D). A small region of interest from (B), that resolves the trimer nanorulers and fiducial structures in its rendered image (E), displays the label IDs (F), and the event groups associated with identified image objects (G). Note that a nanoruler and fiducial marker which were close together have been assigned as being a single object (F-G). Cumulative frequency distribution plots of the two structures boxed in (G), i.e. a trimer nanoruler (H) and a fiducial structure (I). Each distribution has been fit with the theoretical exponential distribution to obtain the mean dark time *τ*_*D*_, and thus its inverse, the qPAINT index associated with the object.

To identify image objects the rendered super-resolution image (Fig. 8) was segmented by a global intensity threshold which encapsulated 80% of the total labelling fraction above background levels. The resulting binary mask exhibited objects that were labelled by assigning unique identifiers (IDs) to connected regions, using a “connected component labelling” approach [55], Fig. 9A. This process was implemented in a PYME analysis workflow (see section 7 “PYME software package”) [49]. The labelled image of objects with object IDs (Fig. 9C,F) was analysed further, determining a range of geometrical properties for each detected object, such as lateral position in the image and total object area (Fig. 9D,G). By relating the image pixel coordinates of an object to the event coordinates of the events, a group of events was identified as being associated with the respective object. Using the frame number in which events occurred, a set of dark times could be determined for each object. The mean dark time was obtained by fitting to the associated dark time histogram as described and the object qPAINT index was determined (Fig. 9H and 9I, see also section 7, “Quantitative Analysis with qPAINT”).

Fig. 10 shows a representative image area (Fig. 10A), identified image objects are coloured according to object area (Fig. 10B) and a corresponding view of objects coloured by qPAINT index (Fig. 10C). It is apparent that thresholding has in some cases led to nearby structures becoming part of the same image object. To measure the qPAINT indices of single nanorulers and single fiducial structures reliably, it was necessary to select only those objects that represented single structures. This was achieved by investigating geometric properties of image objects that could be used to identify single structures only. When plotting the number of events associated with an object against the minor to major axis ratio of the objects, two clusters were identified. One showed a minor to major axis ratio close to 1, corresponding to single fiducial structures, and another cluster showed a minor to major axis ratio closer to 0.4 (Fig. 10D), corresponding to single nanorulers. In addition, the number of events for objects appeared to scale in proportion with the qPAINT index (Fig. 10E).

**Figure 10.**
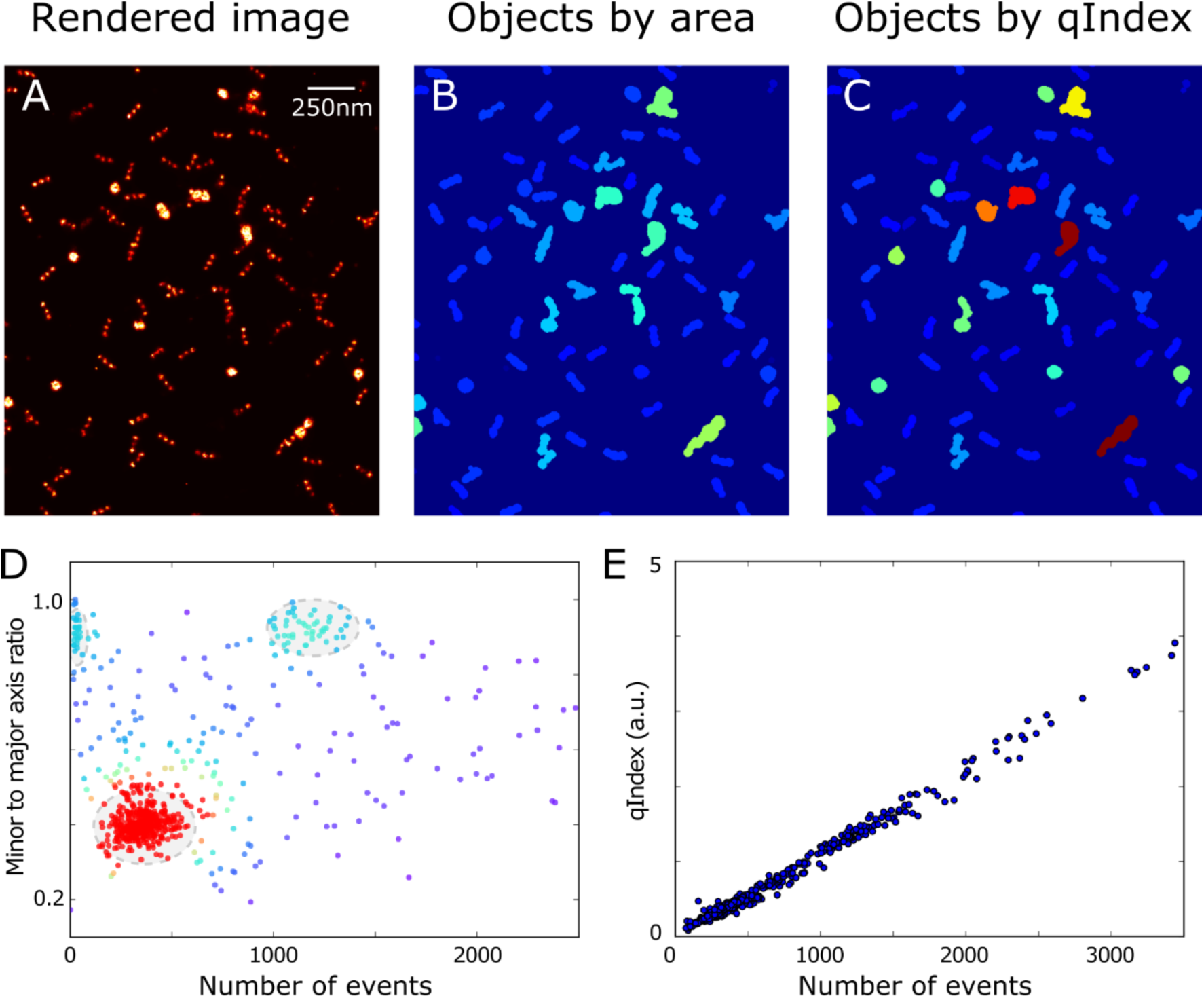
(A) A small region of a rendered super-resolution DNA-PAINT image of a GATTA-PAINT HiRes 40R slide, containing both trimer nanorulers and fiducial marker structures. (B) Connected component labelling of these structures allows them to be filtered by area, clearly revealing overlapping structures. (C) Filtering for qPAINT index highlights the same overlapping structures, but importantly distinguishes between the nanoruler and fiducial markers. (D) Plotting the minor to major axis measurements against qPAINT index for each uniquely identified object produced two dense object clusters. Single nanorulers represent the red cluster, whilst a cluster of the more spherical, fiducial structures made up the second cluster. (E) The qPAINT index, proportional to the number of binding sites, was approximately linearly proportional to the number of detected events.

Fig. 11 illustrates that selecting objects from each of the two clusters identified above yielded objects of single structures, i.e. fiducial structures (Fig. 11A,B) or nanorulers (Fig. 11D,E). This enabled the separate analysis of the two structures in terms of qPAINT analysis by examining their qPAINT indices. As expected, the qPAINT indices fall into a narrow range for each structure that can be approximated by a Gaussian distribution (Fig. 11C,F). We normalised all qPAINT indices by the mean qPAINT index for the group of fiducial structures (Fig. 11C). By comparison, the normalised qPAINT index for the nanorulers was 0.28 (Fig. 11F). The ratio of the two qPAINT indices indicated that there were ∼3.6 times more accessible docking sites in a single fiducial structure than in a single nanoruler. The manufacturer (GATTAquant, private communication) indicated that the ratio of sites, based on the DNA origami designs used, was approximately 4.5, slightly higher than the value we measured. This is compatible with a ∼20% steric hindrance that can be encountered when DNA binding in a dense region of strands [56], as may be occurring with the fiducial marker structure.

**Figure 11.**
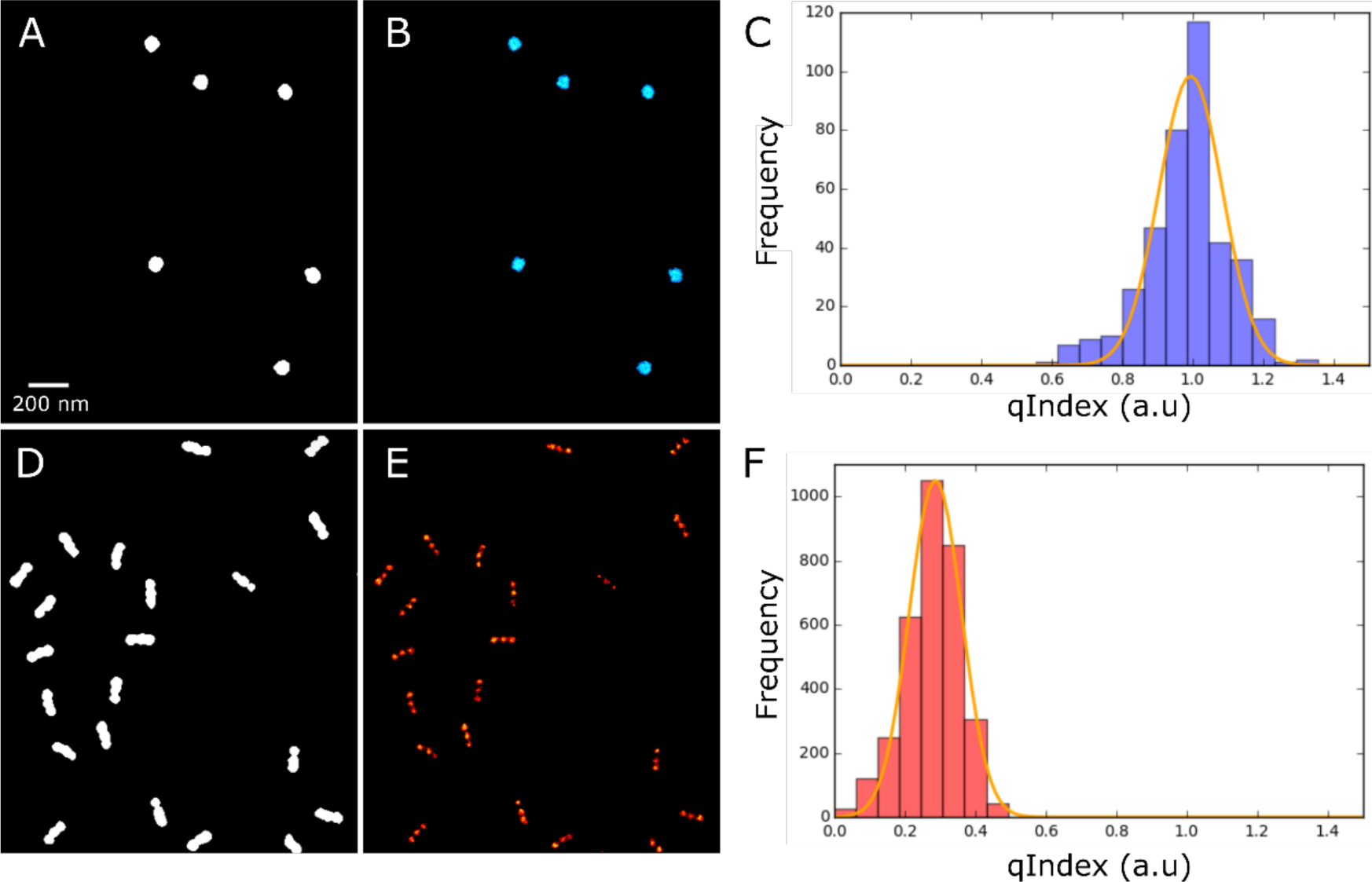
Separate measurements of the qPAINT indices of the fiducial markers and the trimers in the GATTA-PAINT HiRes 40R slide. (A) & (B) Filtered and the rendered super-resolution images of solely the fiducial markers. (C) The measured qPAINT index displayed a single peak which was used to normalise the index. (D)-(F) Repeating the steps for the trimer nanorulers produced a histogram peak at ∼0.28. The ratio between these two values suggested an approximate 3.6 times as many docking sites are available on the fiducial marker as there are present on the nanoruler.

The procedure described above allows the use of this type of slide (and similar test slides) to be used as a convenient test slide for establishing the workflow for qPAINT analysis with a known outcome that can be used to validate against.

## 6. Discussion

We have illustrated the use of 3D test samples to validate the 3D localisation performance of a localisation microscope and demonstrated how to conduct test measurements to validate the qPAINT workflow. When using localisation microscopy the system performance does not only depend on the optical train but also reflects additional aspects such as mechanical stability and drift correction as well as the analysis pipeline used to localise events.

The use of 2D test samples for testing lateral resolution of localisation based super-resolution has already been covered extensively in the literature [25, 29–31, 57]. Our focus on 3D samples and the repurposing of a 2D test sample for establishing qPAINT analysis should help to further establish the use of test samples for this more complex set of SMLM properties and analysis strategies.

While the value of such validation and test measurements should be apparent, it is important to note that imaging synthetic test samples successfully does not prove that a localisation based super-resolution microscope will generate high quality data free from artefacts when imaging often complex biological samples. What it does validate is the imaging and analysis workflow *under the most advantageous conditions* (i.e. bright, long-lasting, well-separated single-molecule events against low backgrounds). If, under these very favourable conditions, the microscope does not perform well or significant artefacts remain, it is almost guaranteed that instrument performance is insufficient to generate meaningful biological data. It is therefore essential that SMLM operators check their system in this manner upon installation, after significant upgrades (this includes software upgrades), setup relocation and any other intervention that could adversely affect system performance.

In fact, when we first used the 3D nanopillar target on our system we only obtained comparably poor data. After troubleshooting the whole acquisition and analysis pipeline a scaling error in the output of our z-tracking feedback loop was eventually identified. After correcting this issue we obtained the much higher quality data shown here. When imaging a biological sample with often unknown 3D structure it can be much more difficult to notice such system performance issues.

A further aspect of the test samples that we use here is the reproducibility of sample properties which can be very helpful during troubleshooting, for example, while investigating unexpected results from imaging a biological sample. The procedures that we describe here can help rule out more obvious types of malfunction of the microscope.

For such test samples to be widely useful they should be readily obtainable and usable by most users of super-resolution microscopes. Two of the samples used here are commercially available and in our hands had good stability over time.

The third sample type is based on custom DNA sequences (that are widely available by custom order from commercial suppliers) and commercially available microspheres. By comparison to directly handling DNA origami for custom made samples (i.e. excluding the cases covered by commercially available ready-made DNA origami test slides used above) the microsphere based samples are much easier to handle for non-experts and assembly of these samples should be accessible to most laboratories working with super-resolution microscopes. While requiring some laboratory skill to make, once made they last several weeks in unsealed open chambers if evaporation is avoided by appropriate storage. One advantage of this preparation is a comparatively low cost per slide. In addition, flexible choice of DNA sequences and dyes is fairly straightforward by ordering appropriately modified DNA.

The use of samples based on DNA-PAINT is very convenient since these samples can often be reused over extended periods, in the case of tightly sealed samples adequate operation over months can be achieved. DNA-PAINT samples are effectively free of photo-bleaching due to the fact that the local diffusion of the solution provides effectively unlimited replenishment of imagers. This means that one can image a sample for a very long period without worrying about the decline of photon yields and even revisit the same region of interest previously imaged, for example, to monitor subtle changes.

This can be very useful in comparing a super-resolution microscope system immediately prior to a move and straight after its relocation to ensure the same or better imaging quality is obtained. We have used some of the samples shown here for exactly this purpose a number of times with good results which helped reduce down-time as we were sufficiently confident to start biological experiments swiftly after relocation.

The use of the DNA-PAINT microspheres and commercial 3D origami samples enable estimation of the resolving power of super-resolution systems in 3D imaging. Imaging modalities are not limited to biplane mode [37] but include other methods, such as astigmatism [58, 59], double-helix [60], phase ramp [39] and multiphase interferometry [61].

The comparatively large microspheres can also be deposited and immobilised into complex biological samples and act as reference objects. The known “cap” structure can be utilised as an indicator to estimate localisation precision and residual sample drift. The large size and high signal-to-noise ratio under brightfield illumination make them also an ideal candidate for sample tracking using correlation methods [62]. In fact, similar microspheres have been used as an effective and affordable way to calibrate the z response of astigmatic 3D localisation [46, 63].

Finally, while we exclusively focus on synthetic test samples here, there is also a very strong case for reproducible biological test samples. We are aware of the cell lines developed by the Ries laboratory as important examples of such an approach [64].

## 7. Materials and Methods

### 7.1 Microsphere Sample Preparation for DNA-PAINT

Lyophilised and high-performance liquid chromatography (HPLC)-purified oligonucleotides (Eurofins Genomics) were resuspended in Tris-EDTA (TE) buffer (pH 8.0, Sigma-Aldrich) at a concentration of 100 µM. For imaging, 500 mM NaCl was added to the buffer. Oligonucleotide sequences were adapted from [65], imager: CTAGATGTAT – 3’Atto 655, docking: 5’Biotin – TTATACATCTA. Microsphere labelling and imaging as described elsewhere [45]. Docking strand oligonucleotides were attached to streptavidin-coated polystyrene colloidal particles (r = 250 nm, binding capacity 500 pmole of biotin per mg of particles, Microparticles GmbH) by suspension of docking strands at 4x excess (1 µM) in TE buffer containing the polystyrene particles (0.5 mg/ml) and 300 mM NaCl. Suspended particles were incubated overnight. Afterwards, unbound oligonucleotides were removed by repeated centrifugation and suspension in TE buffer. Docking-strand coated particles can be stored at 4 °C and used for several days. Non-specific binding of imager strands to the coverslip surface was prevented with PLL-g-PEG (SuSoS). PLL-g-PEG was dissolved in PBS (pH 7.4) at 0.1 mg/ml. Coverslips were coated with PLL-g-PEG, incubated for 2 h, washed with Milli-Q water and dried. The coverslips were attached to a Perspex slide with a circular opening to create an open slide chamber, as shown by Crossman et al. [66]. ∼1 µl of the labelled particle dispersion was dispensed onto the coverslip, partial evaporation of the solution at RT after ∼5 min resulted in stationary particles on the coverslip surface. 100 nm diameter red fluorescent beads (ThermoFisher FluoSpheres F8801, 580 nm peak excitation, 605 nm peak emission) were then deposited into the chamber as fiducial markers for sample drift correction. The fiducial markers were selected so that the dye in these red beads was only weakly excited which helped ensure that the beads were not too bright as compared to the ATTO 655 blinks. At the same time, the “de-tuned” excitation limited the photo-bleaching so that beads produced detectable signals over > 50k frames. The microspheres were imaged in TIRF illumination with an imager concentration of 0.05 nM. Microsphere samples were made according to this recipe in open chambers (see Fig. 2 in [66]) that allow straightforward access to the bathing solution immersing the microspheres.

### 7.2 Experimental Setup

Data were acquired on a Nikon Ti-E inverted microscope with a 60X oil-immersion1.49 NA objective (Nikon Apo TIRF 60X oil) and custom-built optics. Illumination was provided by an arc lamp (Prior Lumen 200S) for calibrating the splitter or a 647nm laser (OMicron LuxX 647-140) for exciting single molecule events. Focussing was provided by a piezo Nanofocusing Z-Drive (Physik Intrumente PD72Z4 400 µm). Sample was held by a slide holder built on a piezo XY stage (Physik Intrumente M-686 Piezoceramic Linear Motors). Fluorescence images were recorded using a sCMOS camera (Andor Zyla 4.2 USB 3.0). For Biplane imaging, an optical splitter (Cairn Research OptoSplit II) was installed between the microscope side port and the camera.

To track lateral sample drift as well as implementing a focus lock during data acquisition, the sample is simultaneously illuminated by the transmitted light from the microscope condenser. Brightfield images were recorded by a separate tracking camera (IDS Imaging UI-3060CP Rev. 2). A band-pass filter (Thorlabs FB450-40) was inserted in the condenser, allowing blue light to propagate to the tracking camera without introducing crosstalk to the fluorescence images. Cross-correlation and interpolation were performed on the brightfield images to estimate the real-time sample drift, which was logged while spooling data for drift correction at the post-processing stage. The measured axial drift was also used to create a negative feedback loop where the position of the objective was locked with respect to the coverslip to maintain focussing [62].

### 7.3 PYME software package

Data acquisition and analysis were implemented in a Python-based software package, Python Microscopy Environment (PYME), which is freely-available at http://python-microscopy.org/. PYME offers customisable acquisition protocols that allow users to predefine a series of hardware setting changes, such as beam intensity or camera settings, at defined times during the acquisition process while providing CPU parallel, real-time analysis. It seamlessly integrates the Python programming language with a robust combination of powerful features for microscopy and is optimised for PALM, STORM and DNA-PAINT super-resolution imaging [49].

### 7.4 Image Acquisition

A 647nm laser was used to excite the ATTO 655 dye in both the GATTA-PAINT and 3D microspheres samples. The excitation beam was adjusted by custom-built 4F optics to TIRF mode with a beam diameter of ∼15 µm and a power density of ∼2kW/cm^2^. Images were captured with an exposure time of 100 ms and a total of 40-70 k frames were acquired in each series.

Point spread functions (PSF) for biplane analysis were obtained by running a Z stack of 200 steps (with an axial step size 50 nm) of sparse sub-resolution (200 nm) dark-red fluorescent beads (ThermoFisher FluoSpheres F8807). A number of such beads are selected, aligned and averaged to extract a 3D PFS which was saved as a TIF file, all using functionality available in PYME.

### 7.5 2D localisation

Detected events in each camera frame were individually isolated with a bandpass filter matched to the expected event size. The centres and peaks of intensity of events were calculated and used as the starting parameters of the fitting procedure. A weighted least squares fit of a Gaussian model, which took into account the non-uniform sCMOS pixel properties, was employed to the unfiltered raw image data with the weights estimated from the Poisson shot noise and Gaussian readout noise of the camera, as described in [31]. Fitting results were obtained by using a Levenburg-Marquardt solver and saved in a data pipeline where post-processing, such as event filtering, was implemented before data visualisation.

### 7.6 3D localisation

In biplane mode 3D localisation, signals from each single molecule event were split into two channels by the splitter and captured simultaneously by the two halves of the camera. With a shift map applied (see section 7.8, “Calibration of Optical Splitter”), the images in the two regions of interest was then combined to form a 3D dataset. In the fitting process an experimentally measured 3D PSF with known axial shift was employed to replace the Gaussian model in 2D localisation. 3D position information was obtained by applying cubic spline interpolation on the measured PSF data. 3D fitting was performed on the raw 3D dataset using a Levenberg-Marquardt algorithm [48, 49]. All of this functionality is implemented in PYME.

To use the PRILM method [39] for 3D localisation, a two-lobed PSF was produced by adding a weak phase ramp that covers half of the objective’s rear pupil plane in the Fourier space. In our case, the phase ramp was produced by utilising a piece of microscope glass slide with an angle of ∼80 arcsec between the two surfaces. With the 60X objective and f = 140 mm tube lens, a separation of ∼500 nm between the two lobes was obtained. The glass slide was installed in a slider which can be inserted underneath the filter turret of the microscope body, allowing convenient switching between conventional 2D and 3D localisation mode. Similar to biplane analysis, in the fitting process cubic spline interpolation was applied to the PSF data and 3D fitting was implemented using the same algorithm, except that event detection was modified to recognise the double-lobed PSF profile. All procedures are implemented in the publicly available PYME software environment which was used both for acquisition and localisation analysis.

### 7.7 Data Visualisation

Filtered localisation clouds were rendered into a super-resolution image using the jittered triangulation method as described in [67] and implemented in PYME. The rendered image was saved as a 16-bit greyscale TIF image with a typical pixel scaling of 5 nm/pixel. The brightness of each pixel was linearly proportional to the local density of localised markers.

### 7.8 Calibration of Optical Splitter

For 3D localisation with the biplane method, it is important to account for differential shifts between the two channels of the optical splitter. Shifts of several tens of nanometers or more are to be expected and typically cannot be corrected merely by precise alignment. Splitter calibration was implemented by using a field of dense 200 nm fluorescent beads (ThermoFisher, dark red FluoSpheres F8807) scanning across the entire region of interest. Illumination was provided by a widefield arclamp to excite as large area as possible. The imaging plane was selected at half way between the two focal planes so that all beads are detectable in both channels. Data acquisition was controlled by a pre-set protocol which automatically moves the sample repeatedly to provide good coverage of the field of view. The positions difference of each bead in the two channels were measured, followed by smoothing spline interpolation which produces a shift vector map with high precision (Fig. 12) [48]. In the fitting process, the vector map is included in the model function which takes one channel as the reference channel and shifts the other channel with respect to it using the measured shift at each lateral position. This is achieved not by shifting the raw data but by modifying the fitting function so that the fit function in channel 2 is offset by the measured amount versus the fit function in channel 1. This procedure is implemented in PYME which uses the shift vector map as input in addition to the PSF when analysing the raw frame data.

**Figure 12.**
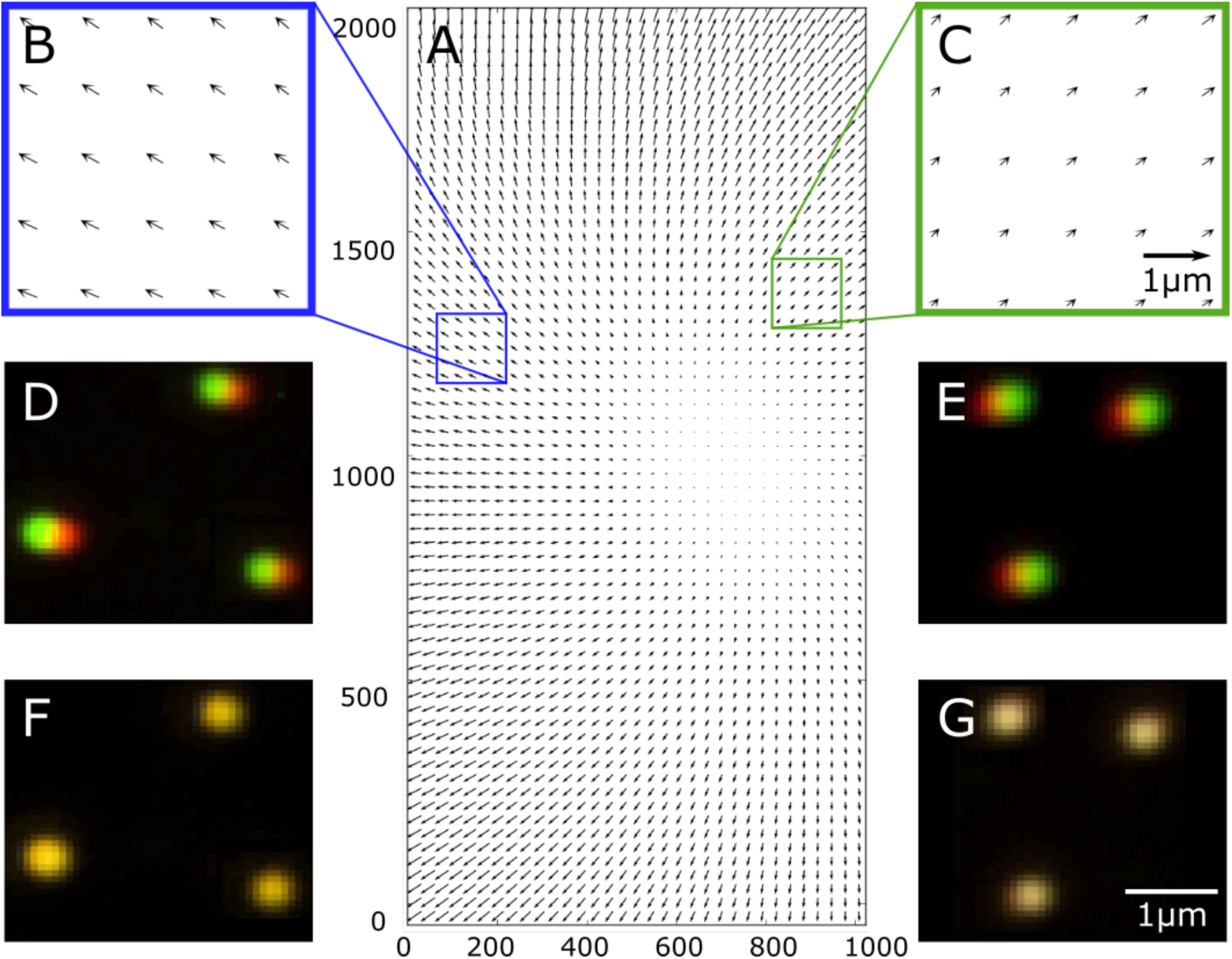
(A) A shift vector map depicting the distortion between the two channels of the optical splitter. (B) & (C) The direction and amount of shift are highly position-dependent and typically cannot be eliminated by precise optical alignment. (D) & (E) Due to the distortion, images of the same object (e.g. sub-resolution beads) are detected at different positions shown as red and green in the two channels. (F) & (G) The distortion is corrected by applying the measured shift map on the corresponding positions, resulting in now overlapping red and green images (yellow).

### 7.9 Axial correction of refractive index mismatch

Refractive index mismatch is an effect commonly encountered when imaging samples that are mounted in a medium whose refractive index (*n*_1_) is different from that of the immersion of the objective (*n*_2_) [52]. Due to the additional refraction at the interface between the mounting medium and the coverslip, light rays emerging out of the front lens of the objective are bent either toward or away from the optical axis, and focus to a plane whose position is determined by the difference of the two refractive indices. The shift of the focal plane leads to a non-constant but z-dependent axial magnification of the optical system. When *n*_1_ < *n*_2_, rays are refracted toward the optical axis, moving the actual focal plane toward the coverslip. In this case, the foreshortened focal length induces an elongation effect on the localisation data cloud which should be corrected by a z foreshortening factor < 1 to reconstruct the true 3D structure of the specimen. When *n*_1_ > *n*_2_, rays are bent away from the optical axis moving the actual focal plane farther away the coverslip, resulting in compression of the cloud with a z factor >1. When *n*_1_ = *n*_2_, the two refractive indices are matched and no correction is required.

Axial foreshortening (*n*_1_ < *n*_2_) is generally inevitable in 3D imaging of DNA-PAINT as the sample is often in aqueous solution whilst oil immersion is used with a high NA objective to enable TIRF or HILO illumination. Measuring the z factor requires use of a sample with known structure. For the 500 nm microsphere, an ellipsoid model with a scalable z axis was used as the fitting model:

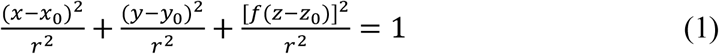

where (*x*_0_, *y*_0_, *z*_0_) is the center of the ellipsoid. *r* is the radius of the equatorial plane and *f* is the z foreshortening factor.

The GATTA-PAINT 3D HiRes 80R DNA origami sample contains nanorulers anchored in various orientations and can also be used as a calibration sample for measuring the z factor. For each nanoruler, a Gaussian mixture model [51] was used to find the centers of its two drift-corrected clouds, from which the measured nanoruler length *L*_*m*_ and angle *θ*_*m*_ can be obtained. This process was repeated for a set of 20 randomly selected nanorulers and the data points (*L*_*m*_, *θ*_*m*_) were then fitted by a model described in [26]:

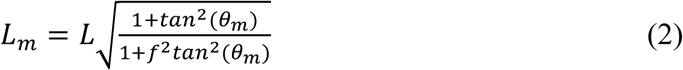

where *L* is the actual length of the nanorulers and *f* is the z foreshortening factor.

Finally, the measured z coordinate *z*_*m*_ of each event was multiplied by the z foreshortening factor to obtain the actual z position:

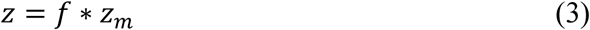

### 7.10 Quantitative Analysis with qPAINT

qPAINT is an analysis approach for DNA-PAINT data that enables quantitative analysis of the target sample without spatially resolving it. In qPAINT, the number of binding sites is assessed by statistical analysis of the temporal properties of the hybridisation between the DNA docking and imager strands. Using the kinetic model for DNA association and dissociation, the mean unbinding time (fluorescence dark time) *τ*_1_ of a single docking site can be expressed as:

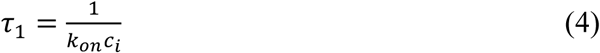

where *k*_*on*_ is the second order on-rate which is constant at a given temperature and buffer makeup, and *c*_*i*_ is the concentration of imager strands in solution.

Considering a target with *N* docking sites, due to the stochastic and reversible binding, the measured mean fluorescence dark time *τ*_*D*_ should be *N* times shorter, i.e.:

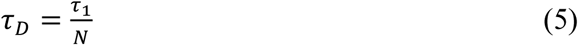

combining the above formulae yields:

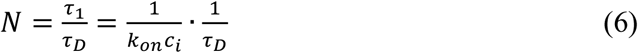

Determination of the absolute number of docking strands generally needs calibration experiments with a structure known to contain only a single or a known number of docking sites [8, 36]. In the absence of such a calibration, the inverse of the dark time, 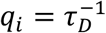, which we call the qPAINT index, can be used as an uncalibrated quantity proportional to the number of docking sites.

It then follows that the ratio *R* of the number of docking sites in two structures can be obtained by the ratio of their respective qPAINT indices:

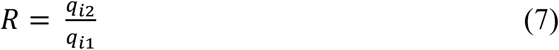

where *q*_*i1*_ and *q*_*i2*_ are the qPAINT indices of structures 1 and 2, respectively. The measurement of the mean dark time for a given structure is illustrated in the main text and shown in Figs. 8 and 9.

## 8. Acknowledgements

This research was supported by grants from Human Frontier Science Program (RGP0027/2013) and the Engineering and Physical Sciences Research Council (EP/N008235/1).

## References

[1] E. Betzig, G.H. Patterson, R. Sougrat, O.W. Lindwasser, S. Olenych, J.S. Bonifacino, M.W. Davidson, J. Lippincott-Schwartz, and H.F. Hess, Imaging intracellular fluorescent proteins at nanometer resolution. Science, 2006. 313(5793): p. 1642–1645.

[2] M.J. Rust, M. Bates, and X. Zhuang, Sub–diffraction–limit imaging by stochastic optical reconstruction microscopy (STORM). Nat Methods, 2006. 3(10): p. 793–5.

[3] R. Jungmann, M.S. Avendano, J.B. Woehrstein, M. Dai, W.M. Shih, and P. Yin, Multiplexed 3D cellular super–resolution imaging with DNA–PAINT and Exchange–PAINT. Nat Methods, 2014. 11(3): p. 313–8.

[4] M. Mikhaylova, B.M. Cloin, K. Finan, R. van den Berg, J. Teeuw, M.M. Kijanka, M. Sokolowski, E.A. Katrukha, M. Maidorn, F. Opazo, S. Moutel, M. Vantard, F. Perez, P.M. van Bergen en Henegouwen, C.C. Hoogenraad, H. Ewers, and L.C. Kapitein, Resolving bundled microtubules using anti–tubulin nanobodies. Nat Commun, 2015. 6: p. 7933.

[5] M. Sauer and M. Heilemann, Single–Molecule Localization Microscopy in Eukaryotes. Chem Rev, 2017. 117(11): p. 7478–7509.

[6] C. Cremer, A. Szczurek, F. Schock, A. Gourram, and U. Birk, Super–resolution microscopy approaches to nuclear nanostructure imaging. Methods, 2017. 123: p. 11–32.

[7] D. Baddeley, I.D. Jayasinghe, L. Lam, S. Rossberger, M.B. Cannell, and C. Soeller, Optical single–channel resolution imaging of the ryanodine receptor distribution in rat cardiac myocytes. Proc Natl Acad Sci U S A, 2009. 106(52): p. 22275–80.

[8] I. Jayasinghe, A.H. Clowsley, R. Lin, T. Lutz, C. Harrison, E. Green, D. Baddeley, L. Di Michele, and C. Soeller, True Molecular Scale Visualization of Variable Clustering Properties of Ryanodine Receptors. Cell Rep, 2018. 22(2): p. 557–567.

[9] S.J. Sahl, S.W. Hell, and S. Jakobs, Fluorescence nanoscopy in cell biology. Nat Rev Mol Cell Biol, 2017. 18(11): p. 685–701.

[10] K.I. Hng and D. Dormann, ConfocalCheck--a software tool for the automated monitoring of confocal microscope performance. PLoS One, 2013. 8(11): p. e79879.

[11] A. Ferrand, K.D. Schleicher, N. Ehrenfeuchter, W. Heusermann, and O. Biehlmaier, Using the NoiSee workflow to measure signal-to-noise ratios of confocal microscopes. Scientific Reports, 2019. 9(1): p. 1165.

[12] A.H. Klemm, A.W. Thomae, K. Wachal, and S. Dietzel, Tracking Microscope Performance: A Workflow to Compare Point Spread Function Evaluations Over Time. Microsc Microanal, 2019: p. 1–6.

[13] R.M. Zucker and O.T. Price, Practical confocal microscopy and the evaluation of system performance. Methods, 1999. 18(4): p. 447–58.

[14] A.D. Corbett, M. Shaw, A. Yacoot, A. Jefferson, L. Schermelleh, T. Wilson, M. Booth, and P.S. Salter, Microscope calibration using laser written fluorescence. Optics Express, 2018. 26(17): p. 21887–21899.

[15] M. Bellec, A. Royon, B. Bousquet, K. Bourhis, M. Treguer, T. Cardinal, M. Richardson, and L. Canioni, Beat the diffraction limit in 3D direct laser writing in photosensitive glass. Opt Express, 2009. 17(12): p. 10304–18.

[16] H. Nyquist, Certain Topics in Telegraph Transmission Theory. Transactions of the American Institute of Electrical Engineers, 1928. 47(2): p. 617–644.

[17] C.E. Shannon, Communication in the Presence of Noise. Proceedings of the IRE, 1949. 37(1): p. 10–21.

[18] J.S. Biteen, M.A. Thompson, N.K. Tselentis, G.R. Bowman, L. Shapiro, and W.E. Moerner, Super-resolution imaging in live Caulobacter crescentus cells using photoswitchable EYFP. Nat Methods, 2008. 5(11): p. 947–9.

[19] J.B. Pawley, Handbook Of Biological Confocal Microscopy. 2006.

[20] K.I. Mortensen, L.S. Churchman, J.A. Spudich, and H. Flyvbjerg, Optimized localization analysis for single-molecule tracking and super-resolution microscopy. Nat Methods, 2010. 7(5): p. 377–81.

[21] R.E. Thompson, D.R. Larson, and W.W. Webb, Precise nanometer localization analysis for individual fluorescent probes. Biophys J, 2002. 82(5): p. 2775–83.

[22] H. Deschout, F. Cella Zanacchi, M. Mlodzianoski, A. Diaspro, J. Bewersdorf, S.T. Hess, and K. Braeckmans, Precisely and accurately localizing single emitters in fluorescence microscopy. Nat Methods, 2014. 11(3): p. 253–66.

[23] Y. Lin, J.J. Long, F. Huang, W.C. Duim, S. Kirschbaum, Y. Zhang, L.K. Schroeder, A.A. Rebane, M.G. Velasco, A. Virrueta, D.W. Moonan, J. Jiao, S.Y. Hernandez, Y. Zhang, and J. Bewersdorf, Quantifying and optimizing single-molecule switching nanoscopy at high speeds. PLoS One, 2015. 10(5): p. e0128135.

[24] A. Small and S. Stahlheber, Fluorophore localization algorithms for super-resolution microscopy. Nat Methods, 2014. 11(3): p. 267–79.

[25] C. Steinhauer, R. Jungmann, T.L. Sobey, F.C. Simmel, and P. Tinnefeld, DNA Origami as a Nanoscopic Ruler for Super-Resolution Microscopy. Angewandte Chemie International Edition, 2009. 48(47): p. 8870–8873.

[26] J.J. Schmied, C. Forthmann, E. Pibiri, B. Lalkens, P. Nickels, T. Liedl, and P. Tinnefeld, DNA Origami Nanopillars as Standards for Three-Dimensional Superresolution Microscopy. Nano Letters, 2013. 13(2): p. 781–785.

[27] C.R. Copeland, J. Geist, C.D. McGray, V.A. Aksyuk, J.A. Liddle, B.R. Ilic, and S.M. Stavis, Subnanometer localization accuracy in widefield optical microscopy. Light: Science & Applications, 2018. 7(1): p. 31.

[28] F.C. Zanacchi, C. Manzo, A.S. Alvarez, N.D. Derr, M.F. Garcia-Parajo, and M. Lakadamyali, A DNA origami platform for quantifying protein copy number in super-resolution. Nat Methods, 2017. 14(8): p. 789–792.

[29] J.J. Schmied, A. Gietl, P. Holzmeister, C. Forthmann, C. Steinhauer, T. Dammeyer, and P. Tinnefeld, Fluorescence and super-resolution standards based on DNA origami. Nat Methods, 2012. 9(12): p. 1133–4.

[30] M. Raab, I. Jusuk, J. Molle, E. Buhr, B. Bodermann, D. Bergmann, H. Bosse, and P. Tinnefeld, Using DNA origami nanorulers as traceable distance measurement standards and nanoscopic benchmark structures. Scientific Reports, 2018. 8(1): p. 1780.

[31] R. Lin, A.H. Clowsley, I.D. Jayasinghe, D. Baddeley, and C. Soeller, Algorithmic corrections for localization microscopy with sCMOS cameras - characterisation of a computationally efficient localization approach. Opt Express, 2017. 25(10): p. 11701–11716.

[32] M. Reuss, F. Fördős, H. Blom, O. Öktem, B. Högberg, and H. Brismar, Measuring true localization accuracy in super resolution microscopy with DNA-origami nanostructures. New Journal of Physics, 2017. 19(2): p. 025013.

[33] G. Ball, J. Demmerle, R. Kaufmann, I. Davis, I.M. Dobbie, and L. Schermelleh, SIMcheck: a Toolbox for Successful Super-resolution Structured Illumination Microscopy. Sci Rep, 2015. 5: p. 15915.

[34] D. Sage, H. Kirshner, T. Pengo, N. Stuurman, J. Min, S. Manley, and M. Unser, Quantitative evaluation of software packages for single-molecule localization microscopy. Nat Methods, 2015. 12(8): p. 717–24.

[35] D. Sage, T.-A. Pham, H. Babcock, T. Lukes, T. Pengo, J. Chao, R. Velmurugan, A. Herbert, A. Agrawal, S. Colabrese, A. Wheeler, A. Archetti, B. Rieger, R. Ober, G.M. Hagen, J.-B. Sibarita, J. Ries, R. Henriques, M. Unser, and S. Holden, Super-resolution fight club: A broad assessment of 2D & 3D single-molecule localization microscopy software. bioRxiv, 2018: p. 362517.

[36] R. Jungmann, M.S. Avendano, M. Dai, J.B. Woehrstein, S.S. Agasti, Z. Feiger, A. Rodal, and P. Yin, Quantitative super-resolution imaging with qPAINT. Nat Methods, 2016. 13(5): p. 439–42.

[37] M.F. Juette, T.J. Gould, M.D. Lessard, M.J. Mlodzianoski, B.S. Nagpure, B.T. Bennett, S.T. Hess, and J. Bewersdorf, Three-dimensional sub-100 nm resolution fluorescence microscopy of thick samples. Nat Methods, 2008. 5(6): p. 527–9.

[38] M.J. Mlodzianoski, M.F. Juette, G.L. Beane, and J. Bewersdorf, Experimental characterization of 3D localization techniques for particle-tracking and super-resolution microscopy. Opt Express, 2009. 17(10): p. 8264–77.

[39] D. Baddeley, M.B. Cannell, and C. Soeller, Three-dimensional sub-100 nm superresolution imaging of biological samples using a phase ramp in the objective pupil. Nano Research, 2011. 4(6): p. 589–598.

[40] A. Sharonov and R.M. Hochstrasser, Wide-field subdiffraction imaging by accumulated binding of diffusing probes. Proc Natl Acad Sci U S A, 2006. 103(50): p. 18911–6.

[41] R. Jungmann, C. Steinhauer, M. Scheible, A. Kuzyk, P. Tinnefeld, and F.C. Simmel, Single-molecule kinetics and superresolution microscopy by fluorescence imaging of transient binding on DNA origami. Nano Lett, 2010. 10(11): p. 4756–61.

[42] D. Axelrod, Cell-substrate contacts illuminated by total internal reflection fluorescence. J Cell Biol, 1981. 89(1): p. 141–5.

[43] M. Tokunaga, N. Imamoto, and K. Sakata-Sogawa, Highly inclined thin illumination enables clear single-molecule imaging in cells. Nat Methods, 2008. 5(2): p. 159–61.

[44] P.W. Rothemund, Folding DNA to create nanoscale shapes and patterns. Nature, 2006. 440(7082): p. 297–302.

[45] T. Lutz, A.H. Clowsley, R. Lin, S. Pagliara, L. Di Michele, and C. Soeller, Versatile multiplexed superresolution imaging of nanostructures by Quencher-Exchange-PAINT. Nano Research, 2018. 11(12): p. 6141–6154.

[46] P. Delcanale, B. Miret-Ontiveros, M. Arista-Romero, S. Pujals, and L. Albertazzi, Nanoscale Mapping Functional Sites on Nanoparticles by Points Accumulation for Imaging in Nanoscale Topography (PAINT). ACS Nano, 2018. 12(8): p. 7629–7637.

[47] C. Cabriel, N. Bourg, G. Dupuis, and S. Leveque-Fort, Aberration-accounting calibration for 3D single-molecule localization microscopy. Opt Lett, 2018. 43(2): p. 174–177.

[48] D. Baddeley, D. Crossman, S. Rossberger, J.E. Cheyne, J.M. Montgomery, I.D. Jayasinghe, C. Cremer, M.B. Cannell, and C. Soeller, 4D super-resolution microscopy with conventional fluorophores and single wavelength excitation in optically thick cells and tissues. PLoS One, 2011. 6(5): p. e20645.

[49] The PYthon Microscopy Environment. [cited 2019 27/01]; Available from: https://python-microscopy.org/.

[50] GATTAquant. product properties of GATTA-PAINT 3D HiRes Nanopillars. [cited 2019 27/01]; Available from: http://www.gattaquant.com/products/localization-based/gatta-paint-3d.html.

[51] I.B. Shabat. What is a Gaussian mixture model (GMM) - 3D point cloud classification primer. [cited 2019 04/03]; Available from: http://www.itzikbs.com/gaussian-mixture-model-gmm-3d-point-cloud-classification-primer.

[52] A. Egner and S.W. Hell, Aberrations in Confocal and Multi-Photon Fluorescence Microscopy Induced by Refractive Index Mismatch, in Handbook Of Biological Confocal Microscopy, J.B. Pawley, Editor. 2006, Springer US: Boston, MA. p. 404–413.

[53] GATTAquant. product properties of GATTA-PAINT HiRes nanorulers. [cited 2019 27/01]; Available from: http://www.gattaquant.com/products/localization-based/gatta-paint-hires.html.

[54] P. Rubin-Delanchy, G.L. Burn, J. Griffie, D.J. Williamson, N.A. Heard, A.P. Cope, and D.M. Owen, Bayesian cluster identification in single-molecule localization microscopy data. Nat Methods, 2015. 12(11): p. 1072–6.

[55] K. Wu, E. Otoo, and A. Shoshani, Optimizing connected component labeling algorithms. Medical Imaging. Vol. 5747. 2005: SPIE. 12.

[56] A.V. Pinheiro, J. Nangreave, S. Jiang, H. Yan, and Y. Liu, Steric Crowding and the Kinetics of DNA Hybridization within a DNA Nanostructure System. ACS Nano, 2012. 6(6): p. 5521–5530.

[57] G. Tortarolo, M. Castello, A. Diaspro, S. Koho, and G. Vicidomini, Evaluating image resolution in stimulated emission depletion microscopy. Optica, 2018. 5(1): p. 32–35.

[58] B. Huang, W. Wang, M. Bates, and X. Zhuang, Three-dimensional super-resolution imaging by stochastic optical reconstruction microscopy. Science, 2008. 319(5864): p. 810–3.

[59] H.P. Kao and A.S. Verkman, Tracking of single fluorescent particles in three dimensions: use of cylindrical optics to encode particle position. Biophys J, 1994. 67(3): p. 1291–300.

[60] S.R.P. Pavani, M.A. Thompson, J.S. Biteen, S.J. Lord, N. Liu, R.J. Twieg, R. Piestun, and W.E. Moerner, Three-dimensional, single-molecule fluorescence imaging beyond the diffraction limit by using a double-helix point spread function. Proceedings of the National Academy of Sciences, 2009. 106(9): p. 2995.

[61] G. Shtengel, J.A. Galbraith, C.G. Galbraith, J. Lippincott-Schwartz, J.M. Gillette, S. Manley, R. Sougrat, C.M. Waterman, P. Kanchanawong, M.W. Davidson, R.D. Fetter, and H.F. Hess, Interferometric fluorescent super-resolution microscopy resolves 3D cellular ultrastructure. Proc Natl Acad Sci U S A, 2009. 106(9): p. 3125–30.

[62] R. McGorty, D. Kamiyama, and B. Huang, Active Microscope Stabilization in Three Dimensions Using Image Correlation. Opt Nanoscopy, 2013. 2(1).

[63] A. Auer, T. Schlichthaerle, J.B. Woehrstein, F. Schueder, M.T. Strauss, H. Grabmayr, and R. Jungmann, Nanometer-scale Multiplexed Super-Resolution Imaging with an Economic 3D-DNA-PAINT Microscope. Chemphyschem, 2018. 19(22): p. 3024–3034.

[64] J.V. Thevathasan, Ulf Matti, Maurice Kahnwald, Sudheer Kumar Peneti, Bianca Nijmeijer, Moritz Kueblbeck, Jan Ellenberg, and Jonas Ries, Nuclear Pores as Universal Reference Standards for Quantitative Microscopy. Biophysical Journal, 2019. 116(3).

[65] J. Schnitzbauer, M.T. Strauss, T. Schlichthaerle, F. Schueder, and R. Jungmann, Superresolution microscopy with DNA-PAINT. Nat Protoc, 2017. 12(6): p. 1198–1228.

[66] D.J. Crossman, Y. Hou, I. Jayasinghe, D. Baddeley, and C. Soeller, Combining confocal and single molecule localisation microscopy: A correlative approach to multi-scale tissue imaging. Methods, 2015. 88: p. 98–108.

[67] D. Baddeley, M.B. Cannell, and C. Soeller, Visualization of localization microscopy data. Microsc Microanal, 2010. 16(1): p. 64–72.

